# Chloroplast quality control pathways are dependent on plastid DNA synthesis and nucleotides provided by cytidine triphosphate synthase two

**DOI:** 10.1101/2020.10.28.360057

**Authors:** Kamran Alamdari, Karen E. Fisher, David W. Welsh, Snigdha Rai, Kyle R. Palos, Andrew D. L. Nelson, Jesse D. Woodson

## Abstract

- Reactive oxygen species (ROS) produced in chloroplasts cause oxidative damage, but also signal to initiate chloroplast quality control pathways, cell death, and gene expression. The mechanisms behind these signals are largely unknown.
- The *Arabidopsis thaliana plastid ferrochelatase two* (*fc2*) mutant produces the ROS singlet oxygen in chloroplasts that activates such signaling pathways. Here we mapped one *fc2* suppressor mutation to *CYTIDINE TRIPHOSPHATE SYNTHASE TWO* (*CTPS2*), which encodes one of five enzymes in *Arabidopsis* necessary for *de novo* cytoplasmic CTP (and dCTP) synthesis.
- The *ctps2* mutation reduces chloroplast transcripts and DNA content without similarly affecting mitochondria. Chloroplast nucleic acid content and singlet oxygen signaling are restored by exogenous feeding of the dCTP precursor deoxycytidine, suggesting *ctps2* blocks signaling by limiting nucleotides for chloroplast genome maintenance.
- An investigation of CTPS orthologs in Brassicaceae showed CTPS2 is a member of an ancient lineage distinct from CTPS3. Complementation studies confirmed this analysis; CTPS3 was unable to compensate for CTPS2 function in providing nucleotides for chloroplast DNA and signaling.
- Our studies link cytoplasmic nucleotide metabolism with chloroplast quality control pathways. Such a connection is achieved by a conserved clade of CTPS enzymes that may have evolved specialized functions in providing nucleotides to specific subcellular compartments.

## Introduction

Chloroplasts are specialized plastid organelles responsible for photosynthesis in plants and algae. Their assembly and maintenance is exceedingly complex with up to ~10% of the plant genome dedicated to their function (van Wijk & Baginsky, 2011). This complexity is due to their semi-autonomous nature. Chloroplasts are derived from a cyanobacterial ancestor that was engulfed by a proto-eukaryotic host about 1.2-1.5 billion years ago. Today, chloroplasts still contain small genomes of ~100 genes essential for their functions. The maintenance and expression of this genome is complicated, mostly requiring nuclear-encoded proteins, but also plastid-encoded ribosomal proteins, rRNAs, tRNAs, and an RNA polymerase (del Campo, 2009). Additionally, the copy number of the genome is highly dynamic and may reach up to 80 copies per organelle, possibly to meet the demands of protein synthesis during early chloroplast biogenesis (Sakamoto & Takami, 2018). Chloroplast nucleic acid content is also dependent on cytoplasmic-produced metabolites, such as NTPs and dNTPs required for RNA and DNA synthesis, respectively (Witte & Herde, 2020). How these metabolites are partitioned within the cell to meet various demands is not well understood. As such, the chloroplast proteome of 2,000-3,000 proteins is a product of multiple cellular compartments that work together for assembly, function, maintenance, and finally, disassembly of these essential organelles.

Another major challenge for chloroplasts is photosynthesis itself, which can produce large amounts of reactive oxygen species (ROS) including singlet oxygen (^1^O_2_) at photosystem II and superoxide and hydrogen peroxide at photosystem I (Asada, 2006). This ROS generation is exacerbated by different stressors including excess light (Triantaphylides *et al.*, 2008), drought (Chan *et al.*, 2016), salinity (Suo *et al.*, 2017), and pathogen attack (Lu & Yao, 2018). While ROS can cause significant damage to DNA, lipids, and proteins, plants use them as signaling molecules to inform the cell about stress (abiotic and biotic) and photo-oxidative damage (de Souza *et al.*, 2017; Dogra & Kim, 2019). These signals, once thought to be a single “plastid factor” (Bradbeer *et al.*, 1979), have turned out to be numerous and complex with over 40 different pathways proposed (Chan *et al.*, 2015). While these pathways remain poorly understood, most involve “retrograde” signals that regulate nuclear gene expression in response to damaged or undeveloped chloroplasts. More recently, researchers have demonstrated chloroplast quality control pathways are also activated and lead to the removal of severely damaged chloroplasts from the cell (Izumi & Nakamura, 2018; Woodson, 2019).

To investigate chloroplast quality control signals, we previously identified genetic suppressors of the *Arabidopsis plastid ferrochelatase 2* (*fc2-1*) mutant defective in plastid heme synthesis (Woodson *et al.*, 2015). Although *fc2-1* mutants are relatively healthy in constant 24h light, they produce large amounts of ^1^O_2_ when grown under diurnal light cycling conditions. In these mutants, the reduced FC2 activity creates a bottleneck in the tetrapyrrole (e.g., chlorophyll and heme) pathway leading to the accumulation of the FC2 substrate protoporphyrin IX (Proto) at dawn. Proto (like other unbound tetrapyrroles) is extremely photosensitizable and rapidly produces ^1^O_2_ in the light. In *fc2-1* chloroplasts, this ^1^O_2_ signals to induce chloroplast degradation and eventually cell death. Such cellular damage is not due solely to ^1^O_2_ toxicity, but also to genetically programmed signals (Wagner *et al.*, 2004; Woodson *et al.*, 2015; Shumbe *et al.*, 2016).

A genetic suppressor screen identified *ferrochelatase two suppressor* (*fts*) mutations that allow *fc2-1* seedlings to survive under diurnal light cycles. One mutation affected *Plant U-BOX 4* (*PUB4*), which encodes an E3 ubiquitin ligase necessary for chloroplast degradation (Woodson *et al.*, 2015). We previously hypothesized PUB4 may direct the ubiquitination of damaged chloroplasts, “marking” them for removal and turnover in the central vacuole. Such a chloroplast quality control mechanism appears to be a parallel to chlorophagy (Kikuchi *et al.*, 2020), where the cellular autophagy machinery is used to turn over chloroplasts damaged by UV (Izumi *et al.*, 2017) or excess light (EL) (Nakamura *et al.*, 2018). Collectively, these pathways may exist to identify and remove severely damaged chloroplasts that cannot be repaired for efficient photosynthesis (Woodson, 2016).

While the downstream components involved in degradation pathways are being uncovered, it is still unclear how ^1^O_2_ could initiate a signal within chloroplasts. Recent work suggests plastid gene expression plays an important role in the process. Three *fts* mutants that suppress chloroplast degradation were shown to affect *PENTATRICOPEPTIDE REPEAT CONTAINING PROTEIN 30* (*PPR30*) or “*Mitochondrial*” *TRANSCRIPTIONAL TERMINATION FACTOR 9* (*mTERF9*), which are hypothesized to encode plastid proteins involved in post-transcriptional gene expression (Alamdari *et al.*, 2020). This suggests specific plastid transcripts may play a direct role in propagating ^1^O_2_ signals.

Here we report the mapping of an additional *fts* mutation, *fts39*, affecting the essential gene *CYTIDINE TRIPOSHATE SYNTHASE 2* (*CTPS2*), which encodes one of five conserved cytoplasmic enzymes necessary for de novo CTP synthesis. In addition to blocking chloroplast degradation and signaling, the *ctps2* mutation leads to a specific reduction in plastid gene expression and DNA copy number. These phenotypes are rescued by exogenous deoxycytidine feeding, suggesting the *ctps2* mutation primarily limits dCTP availability for chloroplast function and stress signaling. An investigation of other CTPS orthologs in Brassicaceae showed CTPS2 is a member of an ancient lineage distinct from the conserved CTPS3 ortholog, which was unable to compensate for CTPS2 function. This indicates that plants may have evolved CTPS enzymes with specific functions in providing organelles with nucleotides. This ability to fine tune primary metabolism for chloroplast function reveals a mechanism whereby the cell can control and balance chloroplast development with chloroplast quality control pathways.

## Experimental Procedures

### Biological material, growth conditions, and treatments

*Arabidopsis thaliana* ecotype *Columbia* (Col-0) was the wild type (wt) line and was used to generate all transgenic constructs. T-DNA line GABI_766H08 (*fc2-1*) from the GABI-Kat collection (Kleinboelting *et al.*, 2012) was described previously (Woodson *et al.*, 2011). Double mutant lines were generated via crossing and confirmed by PCR-based markers (primers listed in Table S1).

Seeds were surface sterilized using 30% liquid bleach (v:v) with 0.04% Triton X-100 (v:v) and plated on Linsmaier and Skoog medium pH 5.7 (Caisson Laboratories North Logan, UT) with 0.6% micropropagation type-1 agar powder. Following stratification, seeds were grown at 22°C with constant or diurnal cycling conditions of ~120 μmol m^−2^ sec^−1^ light.

For Pchlide measurement studies, seedlings were germinated on media with two hours of light, wrapped in aluminum foil, grown in the dark for five days, and harvested under dim green light. For excess light treatments, seedlings were grown for four days in constant dim light (5 μmol m^−2^ sec^−1^ white light at 22°C) and shifted to constant excess (650 μmol m^−2^ sec^−1^ using an LED panel (Hettich, Beverly, MA)) or dim (6 μmol m^−2^ sec^−1^) white light at 4°C. Details about bacterial growth, cloning, and vector construction are listed in Methods S1 and S2.

### Gene expression assays

RNA extraction and Real-time quantitative PCR was performed as previously described (Alamdari *et al.*, 2020). Table S1 lists the primers used. Protein extraction and immunoblotting are described in Methods S3.

### Measuring organelle genome copy number

Genomic DNA was extracted from four day-old seedlings using the DNeasy Plant Pro Kit (Qiagen). Real-time qPCR was performed as previously described (Alamdari *et al.*, 2020), except 1.25 ng DNA was used as template per 10 μl reaction. For each compartment, three primer sets were used as probes (nuclear; *ACTIN2* (Chr. 3), *LHCB1.2* (Chr. 1), *LHCB2.2* (Chr. 2), plastid; *psaJ*, *rbcL*, *clpP,* mitochondria; *ccmf*, *atp1*, *cox1*) (Table S1). Organelle copy numbers were calculated for plastids and mitochondria as average levels of plastid probes/nuclear probes and average levels of mitochondria probes/nuclear probes, respectively.

### Tetrapyrrole measurements, singlet oxygen accumulation, cell death assays

Total chlorophyll was extracted using 100% ethanol from seven-day-old seedlings. Cell debris was pelleted thrice at 12,000 x g for 30 min at 4 °C. Chlorophyll was measured spectrophotometrically in 100 μl volumes in a Biotek Synergy H1 Hybrid Reader and path corrections were calculated according to (Warren, 2008). Chlorophyll levels were normalized per seedling (~ 50-130 per line and counted prior to germination). Pchlide was extracted and measured from whole seedlings as previously described (Woodson *et al.*, 2015). Singlet Oxygen Sensor Green (SOSG, Molecular Probes) was used to assess bulk levels of ^1^O_2_ in vivo as previously described (Alamdari *et al.*, 2020). Trypan blue staining to assess cell death was performed as previously described (Alamdari *et al.*, 2020).

### In vivo protein localization and electron Microscopy

Four to five day-old seedlings were imaged on a Zeiss 880 inverted confocal microscope. To stain nuclei, seedlings were incubated in DAPI (1μg/ml) for 20 minutes and rinsed 2x in PBS before imaging. The excitation/emissions wavelengths used were 458/633 for chlorophyll autofluorescence, 405/470 for DAPI, 514/527 for YFP, and 458/480 for CFP. DAPI, YFP, and CFP were scanned on separate tracks with truncated emissions spectra to prevent signal bleed-over. Resulting images were processed using Zen Blue (Carl Zeiss) software and analyzed in Adobe PhotoShop and ImageJ/Fiji. For TEM, four day-old seedlings were fixed, imaged, and analyzed as previously described (Alamdari *et al.*, 2020).

### Phylogenetic analyses

Homologs were identified using *Arabidopsis thaliana* CTPS1-5 as query in CoGe (Lyons *et al.*, 2008) TBLASTN searches in the genomes of *Arabidopsis lyrata* (id 25868), Capsella rubella (id 16754), *Brassica rapa* (id 24668), *Tarenaya hassleriana* (id 34654), *Gossypium raimondii* (id 52149), and *Amborella trichopoda* (id 19525) using an E-value of 1e-10 (default parameters). Protein sequence corresponding to unique hits in each genome were extracted from CoGe and then aligned using MAFFT v7.017 in Geneious (v11.0.4, Biomatters). Maximum likelihood trees were inferred using RAxML-HPC-PTHREADS-SSE3 v8.2 (Stamatakis, 2014) using the GTR+GAMMA model (-m), 1000 rapid bootstrap replicates (-N, -f a), and a random parsimony seed set to 14853.

### Expression analyses

*CTPS* expression values were obtained by reanalyzing *A. thaliana* and *B. rapa* RNA-seq data (Bio-Projects # PRJNA314076, # PRJNA253868 and # PRJNA185152) associated with flower, ovule, silique, stem, leaf, and root tissues. Reads were mapped to the appropriate genome using RMTA (Peri *et al.*, 2020). Per gene read counts were converted to TPM by dividing by the length of the longest isoform in kilobases, and then dividing this value by the sum of reads per kilobase of all genes dividing by one million. TPM values for *A. trichopoda* were obtained from EvoRepro (Ruprecht *et al.*, 2017). Replicate TPM values for the same tissue were averaged, and then normalized z-scores were calculated and graphed in R.

## Results

### Mapping of the *ferrochelatase two suppressor 39* mutant

*fc2-1* mutants are relatively healthy when grown in constant (24h) light conditions, but they suffer cellular degradation in their cotyledons when grown under 6h light / 18h dark diurnal (light cycling) conditions (Woodson *et al.*, 2015). We assessed this cellular degradation by observing the bleaching of cotyledons (Fig. 1a), staining with trypan blue for cell death (Figs. 1b, S1a), and measuring decreased total chlorophyll levels (Fig. 1c). Here we report the characterization of *fts39*, which suppresses and reverses the *fc2-1* cell death phenotype (Fig. 1a). When grown under light cycling conditions, *fc2-1 fts39* seedlings have decreased trypan blue staining (Figs. 1b and S1a) and increased total chlorophyll levels (Fig. 1c) compared to *fc2-1*. Interestingly, *fc2-1 fts39* seedlings were delayed in their germination compared to wt and *fc2-1* after two days in 24h light (Fig. 1d), but recovered after three days (Fig. S1b).

**Figure 1.**
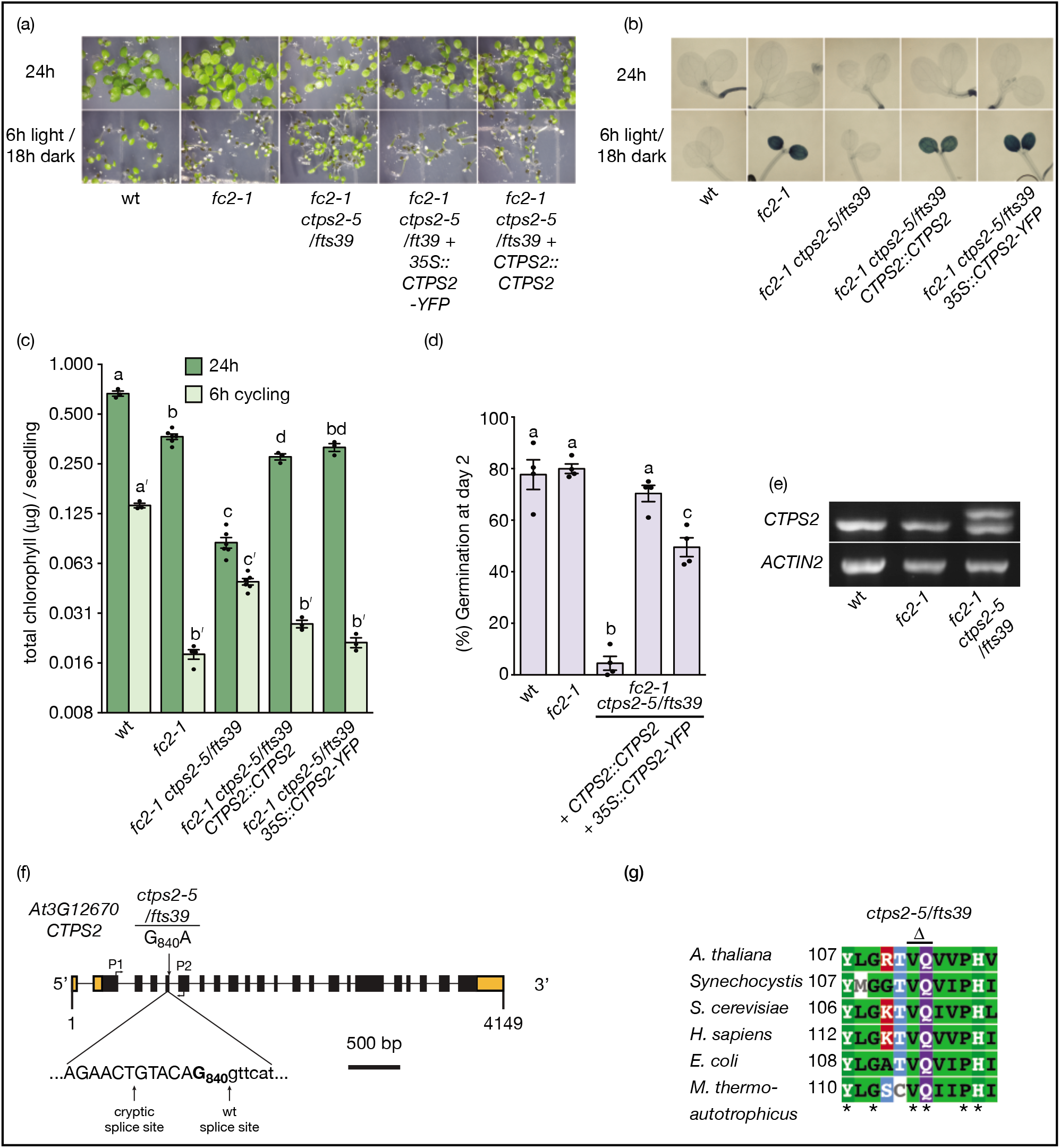
A loss of function *ctps2* allele suppresses cell death in the *fc2-1* mutant. An analysis of *fc2-1 fts39*/*ctps2-5* phenotypes. **A)** Shown are six day-old seedlings grown under constant (24h) light or 6h light / 18h dark diurnal cycling conditions. **B)** Shown are the same seedlings stained with trypan blue. The deep blue color is indicative of cell death. **C)** Shown are mean total chlorophyll levels of six day-old seedlings grown in either constant (24h) light or 6h light / 18 hours dark diurnal cycling (6h cycling) (n ≥ 3 biological replicates) +/- SE. **D)** Mean % germination of seedlings after two days post stratification (n = 4 sets of 50-100 seeds) +/- SE. Germination was defined as cotyledon emergence from the seed coat. Statistical analyses were performed using one-way ANOVA tests and different letters above bars indicate significant differences within data sets determined by Tukey-Kramer post-tests (p-value ≤ 0.05). For C, separate statistical analyses were performed for the different light treatments and the significance for the 6h light / 18h dark diurnal cycling values is denoted by letters with a ʹ. In all bar graphs, closed circles represent individual data points. **E)** DNA gel showing splicing defects of *CTPS2* transcripts in the *fc2-1 fts39/ctps2-5* mutant. cDNA from whole four day-old seedlings was used as template for the PCR. Primers flanking the affected splice site were used for amplification (Table S1). **F)** Schematic of the *CTPS2* (*At3G12670*) gene. Black and orange rectangles indicate the exons and untranslated regions (UTR’s) of the cDNA, respectively. Shown is the position of the single base polymorphism found in the *fts39/ctps2-5* mutant and the resulting cryptic splice site. Also shown are the annealing sites for the primers (P1 (JP1005) and P2 (WLO1400)) used in E. **G)** Protein alignment of various CTPS enzymes across multiple kingdoms and domains. The alignment focuses on 12 amino acids surrounding the area with the two amino acid deletion (V112, Q113) caused by the cryptic splice site in the *ctps2-5* allele.

The causative mutation in *fts39* was previously mapped to a region between 3.82-4.24 mb on the 5’ arm of chromosome three with six single nucleotide polymorphisms (SNPs) (Table S3) (Woodson *et al.*, 2015). To test if any of these SNPs were causing the *fts* phenotype, we cloned each gene (genomic coding region, 5’ UTR, and 3’ UTR) and tested its ability to restore the cell death phenotype of *fc2-1 fts39* mutants. Only the *CTPS2* construct (*AT3G12670*) restored cell death and lower chlorophyll levels in light cycling conditions (Figs. 1a, b, S1a) (Table S4). Furthermore, in 24h light, *CTPS2* restored germination rates (Fig. 1d) and partly restored chlorophyll levels (Figs. 1c). Similar results were observed using a *35S*::*CTPS2-YFP-HA* construct (Figs. 1a-d and S1a and b) that constitutively produced CTPS2 with a fluorescent epitope tag.

*CTPS2* encodes one of five cytoplasmic *Arabidopsis* enzymes predicted to have CTP synthetase activity (Daumann *et al.*, 2018). Using ATP and glutamine, these enzymes catalyze the interconversion of UTP to CTP, which is the rate-limiting step of de novo CTP synthesis (Daumann *et al.*, 2018). Previous reports show *CTPS2* is an essential gene; four complete null *ctps2* alleles caused by T-DNA insertions lead to embryo lethal phenotypes (Table S5) (Daumann *et al.*, 2018; Meinke, 2020). Therefore, the *fts39* (now referred to as *ctps2-5*) mutation is likely a weak allele.

In *ctps2-5*, *CTPS2* carries a conserved G to A point mutation immediately 5’ of intron four. To test if this SNP affects the splicing of the *CTPS2* transcript, we designed primers flanking introns 1-4 that amplify a 368 bp fragment of the wt cDNA. Although wt and *fc2-1* cDNA template led to the amplification of one single PCR product, *fc2-1 ctps2-5* cDNA resulted in two PCR products of similar intensity (Fig. 1e). We cloned each fragment and confirmed their sizes again by PCR (Fig. S1c). Sequencing of these fragments showed the large one contained the wt sequence with correct splicing, while the smaller one was missing six bases immediately 5’ of intron four (bases 331-336 of the coding sequence). Therefore, the *ctps2-5* mutation leads to a cryptic splice site that removes two codons and amino acids (V112 and Q113) from the cDNA and protein, respectively (Fig. 1f).

Comparisons of CTPS homologs from bacteria, archaea, yeast, and humans show these two amino acids are highly conserved (Fig. 1g). A structural analysis of the *E. coli* CTPS protein predicts these two residues are within the active site and the glutamine residue makes contact with the ribose moiety of the UTP substrate (Endrizzi *et al.*, 2004). Although the analysis of total *CTPS2* transcript levels by RT-qPCR indicates they are not significantly changed by the *ctps2-5* mutation (Fig. S1d), we hypothesize approximately half of the CTPS protein is lacking these two conserved residues in this background. To test the possibility that such a protein may have a dominant effect, we overexpressed the *ctps2-5* allele in the *fc2-1* background using the *35S* promoter and a YFP-HA C-terminal fusion. This construct did not produce an *fts* phenotype (nine lines tested) even though significant levels of protein could be detected by immunoblot analysis (Figs. S1e, f). Therefore, *ctps2-5* is a weak recessive allele causing a reduction of functional CTPS2 enzyme.

### The *ctps2-5* mutation blocks stress signals to the nucleus

When suffering from chloroplast photo-oxidative stress under cycling light conditions, *fc2-1* mutants transmit retrograde signals to the nucleus inducing the expression of hundreds of genes (Woodson *et al.*, 2015). To determine if the *ctps2-5* mutation might affect these signals, we used RT-qPCR to monitor expression of six retrograde marker genes for damaged chloroplasts (Woodson *et al.*, 2015), chloroplast ^1^O_2_ (op den Camp *et al.*, 2003), and general stress (Baruah *et al.*, 2009) in four day-old seedlings grown under light cycling conditions. As expected, the expression of all six marker genes was significantly increased one hour after subjective dawn in *fc2-1* seedlings compared to wt (Fig. 2a). In the *fc2-1 ctps2-5* mutant, however, only *ATPase* (*At3g28580*, a marker for ^1^O_2_ chloroplast stress) was significantly induced.

**Figure 2.**
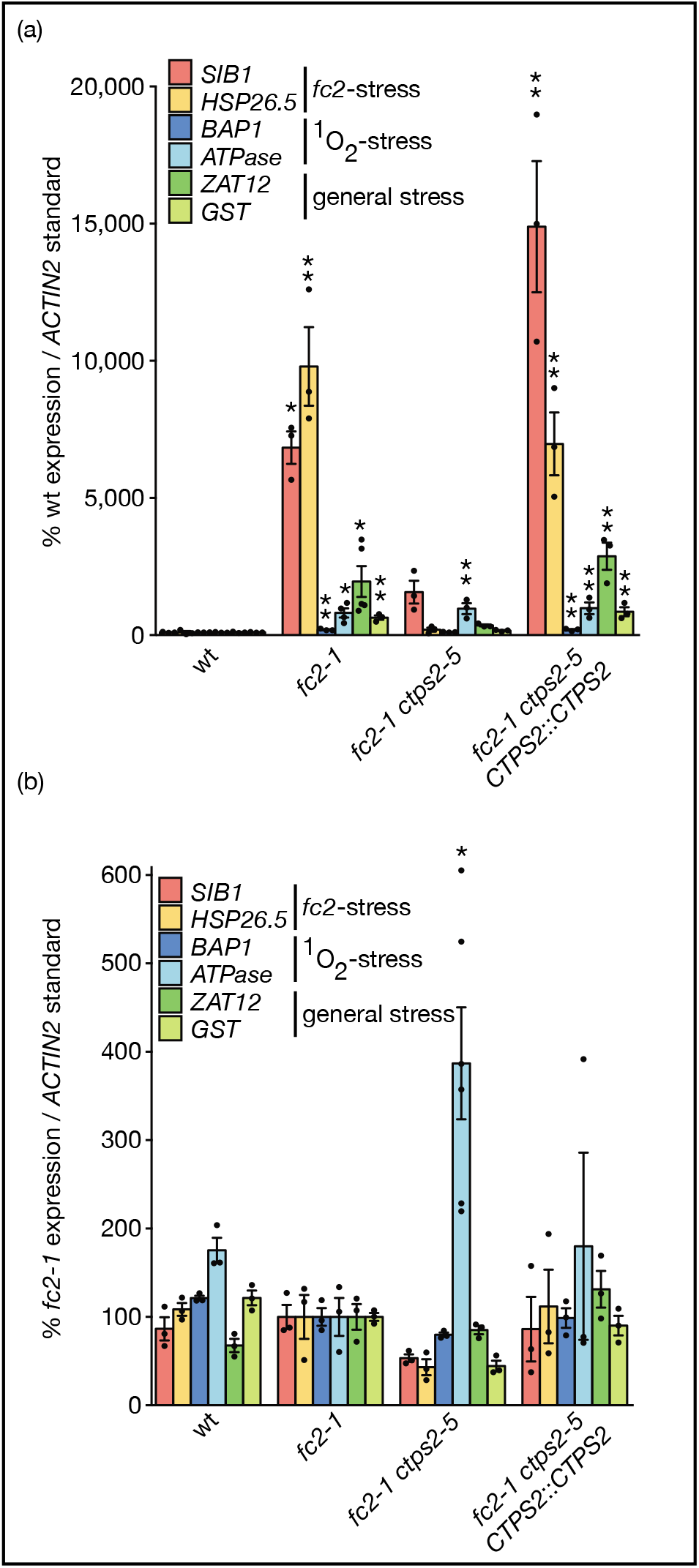
The *ctps2-5* mutation blocks chloroplast stress signaling. An analysis of oxidative stress marker gene transcripts in the *fc2-1 ctps2* mutant. Shown are RT-qPCR analysis of transcripts from four day-old seedlings grown in **A)** 6 hours of light / 18 hours dark diurnal cycling and **B)** constant 24h light conditions. Seedlings were harvested one hour post subjective dawn. Shown are means of biological replicates (n ≥ 3) +/- SE. Statistical analyses were performed by a one-way ANOVA test followed by Dunnett’s multiple comparisons test with the wt (A) or *fc2-1* (B) sample. * and ** indicates an adjusted p value of ≤ 0.05 and ≤ 0.01, respectively. In all bar graphs, closed circles represent individual data points.

Next, we tested 24h light conditions where *fc2-1* mutants do not suffer wholesale chloroplast degradation and cell death. As expected, no single gene was significantly induced in *fc2-1* relative to wt seedlings (Fig. 2b). In *fc2-1 ctps2-5* mutants, however, *ATPase* was induced ~4-fold, *GST* transcripts were significantly reduced, and the other four were unchanged compared to the *fc2-1* single mutant background. Complementation of *fc2-1 ctps2-5* with a wt copy of *CTPS2* was able to fully complement levels of all six transcripts back to *fc2-1* levels in both light conditions. Together, these results indicate *ctps2-5* suppresses the retrograde stress signals in the *fc2-1* mutant.

### *The ctps2-5* mutation leads to delayed chloroplast development

One commonality among *fts* mutants is impairment of chloroplast function and/or development (Woodson *et al.*, 2015). To determine if this is true for the *ctsp2-5* mutant, we monitored the expression of four *Photosynthesis Associated Nuclear Genes* (*PhANGs*) that are repressed by retrograde signals from undeveloped chloroplasts (Woodson *et al.*, 2013). The expression of *LHCB1.2*, *LHCB2.2*, and *CA1* (but not *RCBS2b*) were significantly reduced (relative to *fc2-1*) in four day-old *fc2-1 ctps2-5* seedlings grown in 24h light (Fig. 3a). Complementation of *fc2-1 ctps2-5* with a wt copy of *CTPS2* fully restored transcripts levels of *LHCB1.2* and *CA1. LHCB2.2* transcripts, while not restored to *fc2-1* levels, were significantly increased compared to *fc2-1 ctps2-5* (p-value = 0.001; student’s t-test).

**Figure 3.**
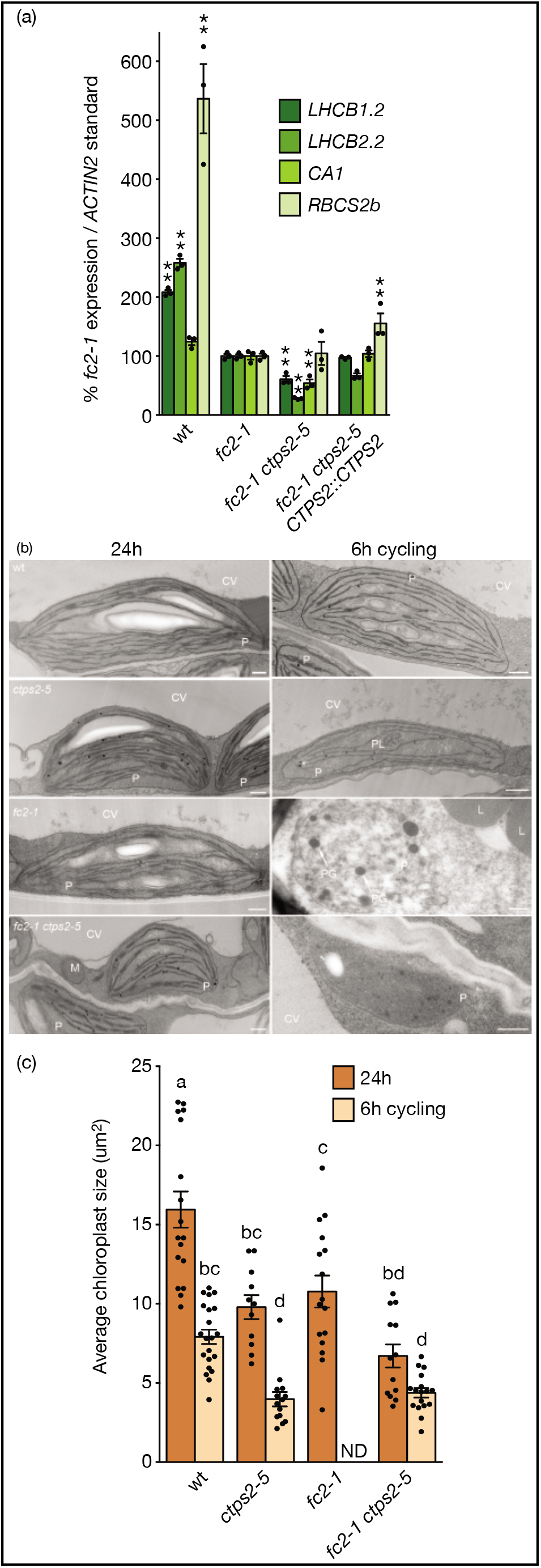
Loss of *CTPS2* leads to impaired chloroplast development and blocks chloroplast degradation. Chloroplast development and degradation was assessed in *fc2-1 ctps2-5* mutants. **A)** Shown are means of expression levels of four PhANGs from four day-old seedlings grown in constant light as measured by RT-qPCR (n = 3 biological replicates) +/- SE. Statistical analysis was performed by a one-way ANOVA test followed by Dunnett’s multiple comparisons test with the *fc2-1* sample. * and ** indicate an adjusted p-value of ≤ 0.05 and ≤ 0.01, respectively. **B)** Shown are representative TEM micrographs of chloroplasts in cotyledon mesophyll cells of four day-old seedlings grown in 24h constant light or 6h light/18h dark diurnal cycling (6h cycling) conditions one hour after subjective dawn. Plastids (P), Nuclei (N), central vacuoles, (CV), lipid bodies (L), plastoglobules (PG), and mitochondria (M) are noted. Scale bars = 500 nm. **C)** Shown are means of the average area (um^2^) of chloroplasts (n ≥ 11 cells) determined from TEM images +/- SE. Chloroplast area in *fc2-1* seedlings grown in 6h cycling light conditions was not determined due to extensive cellular degradation. Statistical analysis was performed by a one-way ANOVA test and different letters above bars indicate significant differences determined by a Tukey-Kramer post-test (p-value ≤ 0.05). In all bar graphs, closed circles represent individual data points.

Next, we visualized the ultrastructure of cotyledon mesophyll cell chloroplasts by transmission electron microscopy (TEM). Consistent with the reduced *PhANG* expression in 24h light, *fc2-1 ctps2-5* chloroplasts were smaller and contained fewer internal membranes than *fc2-1* chloroplasts (Figs. 3b and c). This effect was not unique to the *fc2-1* background as single *ctps2-5* mutants also had smaller chloroplasts compared to wt. Total chloroplast compartment size, however, was not significantly affected by the *ctps2-5* mutation (Fig. S2). Although *fc2-1 ctps2-5* chloroplasts appeared undeveloped, they completely avoided degradation under light cycling conditions (Fig. 3b). Conversely, chloroplasts in *fc2-1* seedlings were often degraded, having poorly defined envelopes, unorganized thylakoid structures, and large plastoglobuli. Together, these results indicate the *ctps2-5* mutation leads to chloroplasts that avoid degradation, but have impaired development.

### The *ctps2-5* mutation uncouples ^1^O_2_ levels from cellular degradation in the *fc2-1* mutant

Under diurnal cycling conditions, *fc2-1* mutants accumulate ^1^O_2_ due to uncontrolled synthesis of Proto or other free tetrapyrroles (Woodson *et al.*, 2015). To monitor flux through this pathway, we measured the accumulation of Pchlide (a precursor of chlorophyll A) in etiolated (dark grown) seedlings. As expected (Scharfenberg *et al.*, 2015; Woodson *et al.*, 2015), the *fc2-1* mutant accumulated significantly more (~2-fold) Pchlide than wt (Fig. 4a). Addition of the *ctps2-5* mutation did not significantly alter Pchlide levels, however, suggesting total flux through the tetrapyrrole pathway was unaltered.

**Figure 4.**
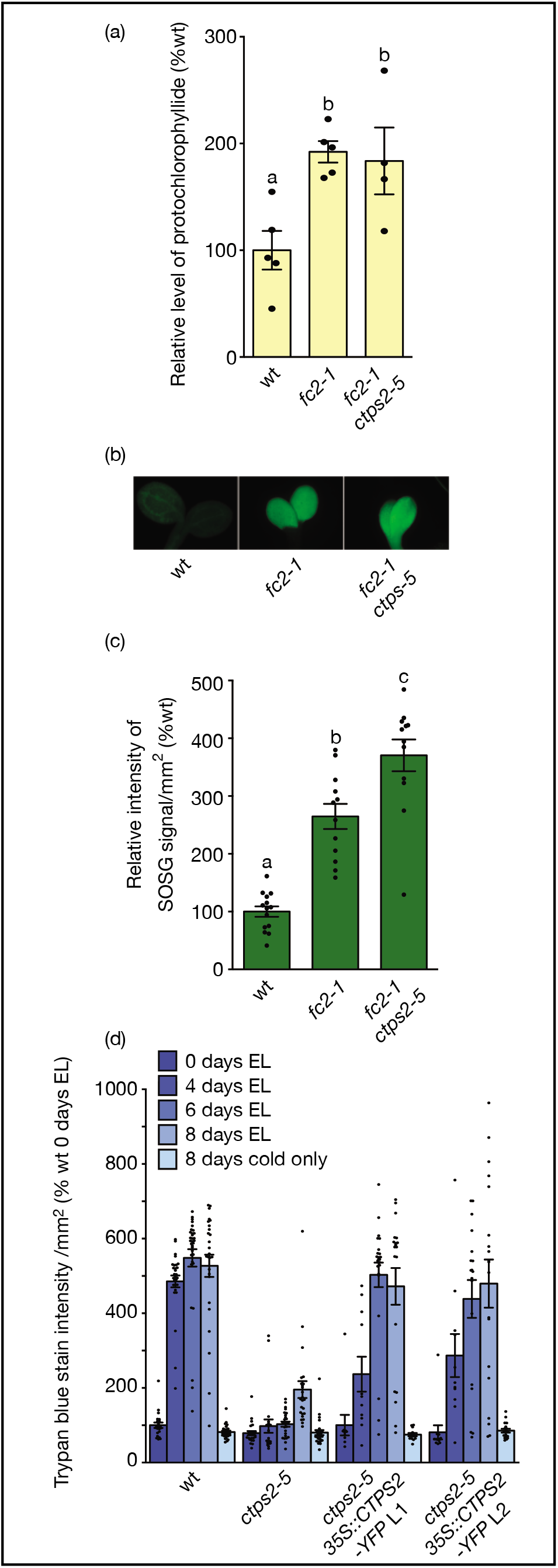
The *ctps2-5* mutation uncouples singlet oxygen levels from chloroplast signaling. The *ctps2-5* mutation does not reduce Pchlide accumulation or ^1^O_2_ levels. **A)** Pchlide levels of five day-old etiolated seedlings grown in constant dark. Shown are means of biological replicate measurements (n ≥ 4) +/- SE. **B** and **C)** ^1^O_2_ levels as measured by SOSG fluorescence in four day-old seedlings grown in diurnal cycling light (6 hours light/18h dark) two hours after subjective dawn. Shown are representative seedlings and means of average intensities of SOSG signal across one cotyledon per seedling (n ≥ 12 biological replicates) +/- SE. **D)** Mean intensities of trypan blue staining in seedlings treated with excess light (EL) and/or 4°C for the indicated amount of time (n ≥ 10 seedlings) +/- SE. Statistical analyses were performed by one-way ANOVA tests and different letters above bars indicate significant differences determined by a Tukey-Kramer post-test (p-value ≤ 0.05). In all bar graphs, closed circles represent individual data points.

Next, we tested if *ctps2-5* was affecting ^1^O_2_ production by using SOSG, a highly sensitive and specific fluorescent probe for ^1^O_2_ (Molecular_Probes, 2004; Flors *et al.*, 2006). Compared to wt, four day-old *fc2-1* mutants grown in cycling light conditions accumulated significantly higher levels of SOSG fluorescence two hours after subjective dawn (Figs. 4b and c). The *ctps2-5* mutation did not suppress this. Instead, *fc2-1 ctsp2-5* accumulated significantly higher levels of SOSG fluorescence compared to *fc2-1*. This result is consistent with the high Pchlide levels in *fc2-1 ctps2-5* mutants, suggesting the *ctps2-5* mutation is affecting the transmission of the ^1^O_2_ signal, rather than the accumulation of the ROS itself.

To determine if the *ctps2-5* mutation could also protect cells from an alternative source of photo-oxidative stress, we exposed single *ctps2-5* mutant seedlings (in a wt *FC2* background) to a combination of excess light (650 μmol m^−2^ sec^−1^) and low temperatures 4°C, which has previously been shown to lead to ^1^O_2_–induced cell death (Meskauskiene *et al.*, 2001; Triantaphylides *et al.*, 2008). Under these conditions, wt seedlings experienced EL-dependent cell death within four days while the *ctps2-5* mutant experienced significantly less cell death for up to eight days (Fig. 4d). A 35S::CTPS2-YFP construct complemented this EL resistance. Together, this suggests the *ctps2-5* mutation also reduces cell death in response to naturally induced photo-oxidative stress.

### *ctps2-5* mutants are defective in plastid gene expression and genome copy number

As plastids and mitochondria must import ribonucleotides for RNA synthesis (Witte & Herde, 2020), we tested if the *ctps2-5* mutation affected plastid gene expression. We performed RT-qPCR to assess steady state transcript levels of eight genes transcribed by the plastid-encoded polymerase (PEP) (*psbA, psbB, psaJ, rbcL, trnEYD*), the nuclear-encoded polymerase (NEP) (*rpoB* and *accD*), or both (*clpP*) (Yagi & Shiina, 2014) by RT-qPCR analysis. Compared to *fc2-1*, the levels of these transcripts (excluding *trnEYD* and *rpoB*) were significantly reduced in *fc2-1 ctps2-5* in 24h light (Fig. 5a). To check the specificity of *ctps2-5* on plastid gene expression, we also tested the expression of six mitochondrial genes, each encoding protein components of six different multi-protein complexes (Fig. S3). The *ctps2-5* mutation did not significantly reduce the expression of any single mitochondrial transcript (p-value ≥ 0.60).

**Figure 5.**
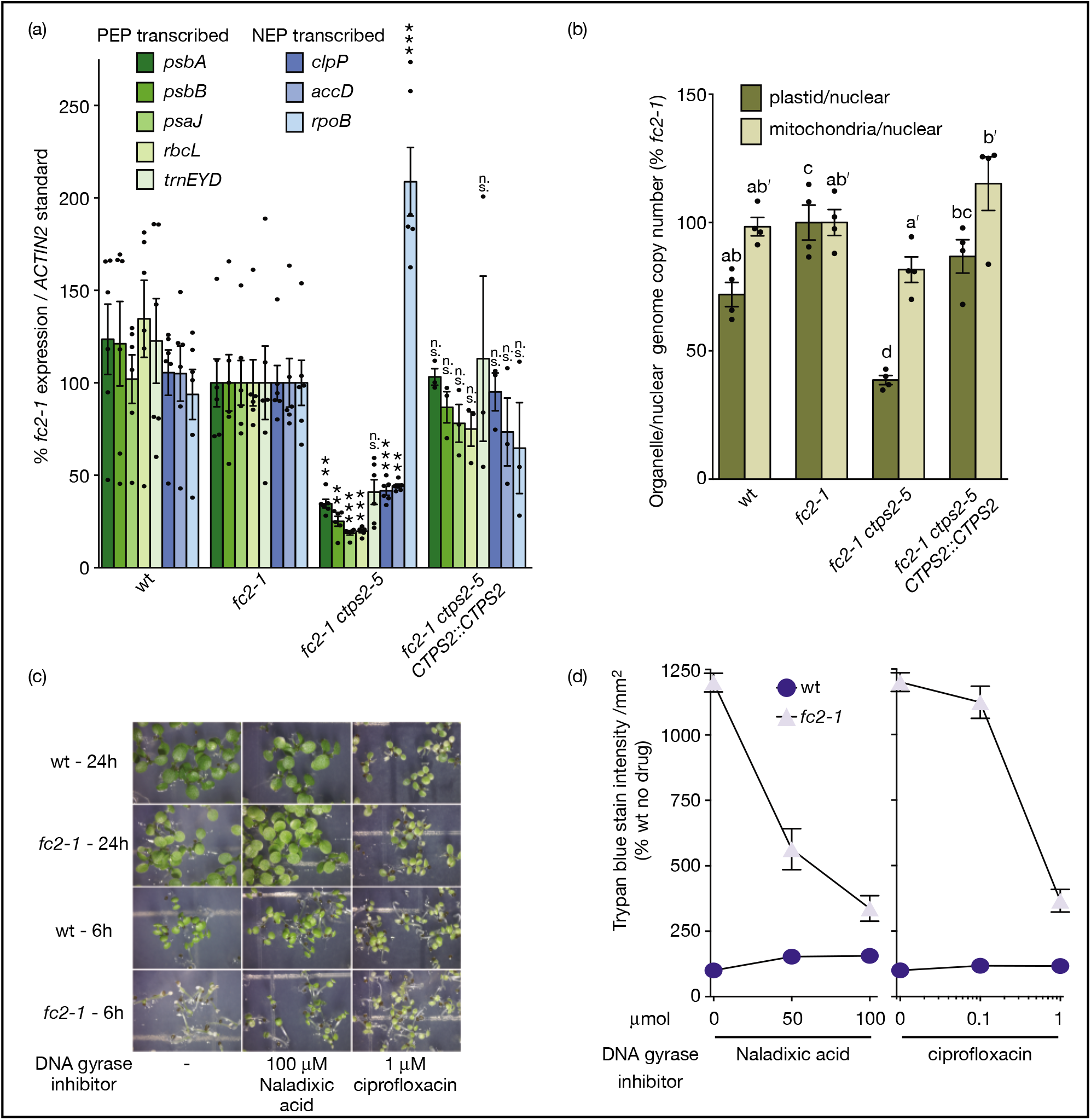
Loss of *CTPS2* leads to a specific reduction in plastid gene expression and genome copy number. An analysis of organelle gene expression and genome copy number in the *fc2-1 ctps2-5* mutant. **A)** Shown are mean levels of plastid transcripts from four day-old seedlings grown in constant light as measured by RT-qPCR (n = 6 biological replicates) +/- SE. Statistical analysis was performed by a one-way ANOVA test followed by Dunnett’s multiple comparisons test with the *fc2-1* sample. ** and *** indicate an adjusted p-value of ≤ 0.01 and ≤ 0.001, respectively. **B)** Shown are the means of relative plastid and mitochondrial genome copy numbers from four day-old seedlings grown in constant light (n = 4 biological replicates) +/- SE. Values were determined by analyzing genomic DNA levels by RT-qPCR using marker probe sets for plastid, mitochondrial, and nuclear genomes (three sets each). Statistical analyses of B was performed using one-way ANOVA tests. Different letters above bars indicate significant differences within data sets determined by a Tukey-Kramer post-test (p-value ≤ 0.05). Separate statistical analyses were performed on plastid and mitochondrial genome copy numbers and the significance for the mitochondrial genome copy number values is denoted by letters with a ʹ. In all bar graphs, closed circles represent individual data points. **C)** Shown are six day-old seedlings grown with the indicated DNA gyrase inhibitor and **D)** the mean intensities of their staining by trypan blue (n ≥ 10 seedlings) +/- SE.

As de novo cytoplasmic CTP synthesis also supplies CDP for dCTP synthesis (Witte & Herde, 2020), we tested whether the *ctps2-5* mutation affected plastid DNA replication and/or genome maintenance. Genomic DNA from four day-old seedlings grown in 24h light was extracted and organelle DNA content was measured by RT-qPCR. Compared to *fc2-1*, the plastid/nuclear genome copy number was reduced over 2-fold in *fc2-1 ctps2-5* mutants, while the mitochondrial/nuclear genome copy number was not significantly reduced (Fig. 5b). Complementation of *fc2-1 ctps2-5* with a wt copy of *CTPS2*::*CTPS2* restored normal expression of plastid transcripts and plastid genome copy number (Figs. 5a, b). Together, these results suggest the *ctps2-5* mutation has a specific effect on plastid gene expression and genome copy number, disproportional to any effect on mitochondrial function.

To test if reduced plastid DNA copy number itself can block cell death in the *fc2-*1 mutant, we grew seedlings in the presence of the prokaryotic DNA gyrase inhibitors Nalidixic acid and ciprofloxacin, which block plastid DNA replication (Gray *et al.*, 2003; Evans-Roberts *et al.*, 2016). The addition of 100 μM Nalidixic acid or 1 μM ciprofloxacin was clearly able to restore growth and block cell death (Figs. 5c and d) of *fc2-1* in 6h cycling light conditions. Together, these results suggest reduced plastid DNA levels are sufficient to suppress the *fc2-1* cell death phenotype.

### *ctps2-5* chloroplasts are limited for dCTP

The reduced plastid genome copy number in *fc2-1 ctps2-5* seedlings suggests their chloroplasts may be limited for dCTP for DNA synthesis. To test this, we supplemented the growth media with 2mM dNTP’s, which has been shown to rescue the phenotypes of nucleotide metabolism mutants (Xu *et al.*, 2020). The addition of dCTP to the media rescued the effect of the *ctps2-5* mutation and led to increased cell death of the *fc2-1 ctps2-5* mutant under cycling light conditions (Figs. 6a, b). dCTP was not toxic in itself, as it had no effect on wt in cycling light or on *fc2-1 ctps2-5* mutants under 24h light. Supplementing media with the other three dNTP’s had no significant effect on cell death.

**Figure 6.**
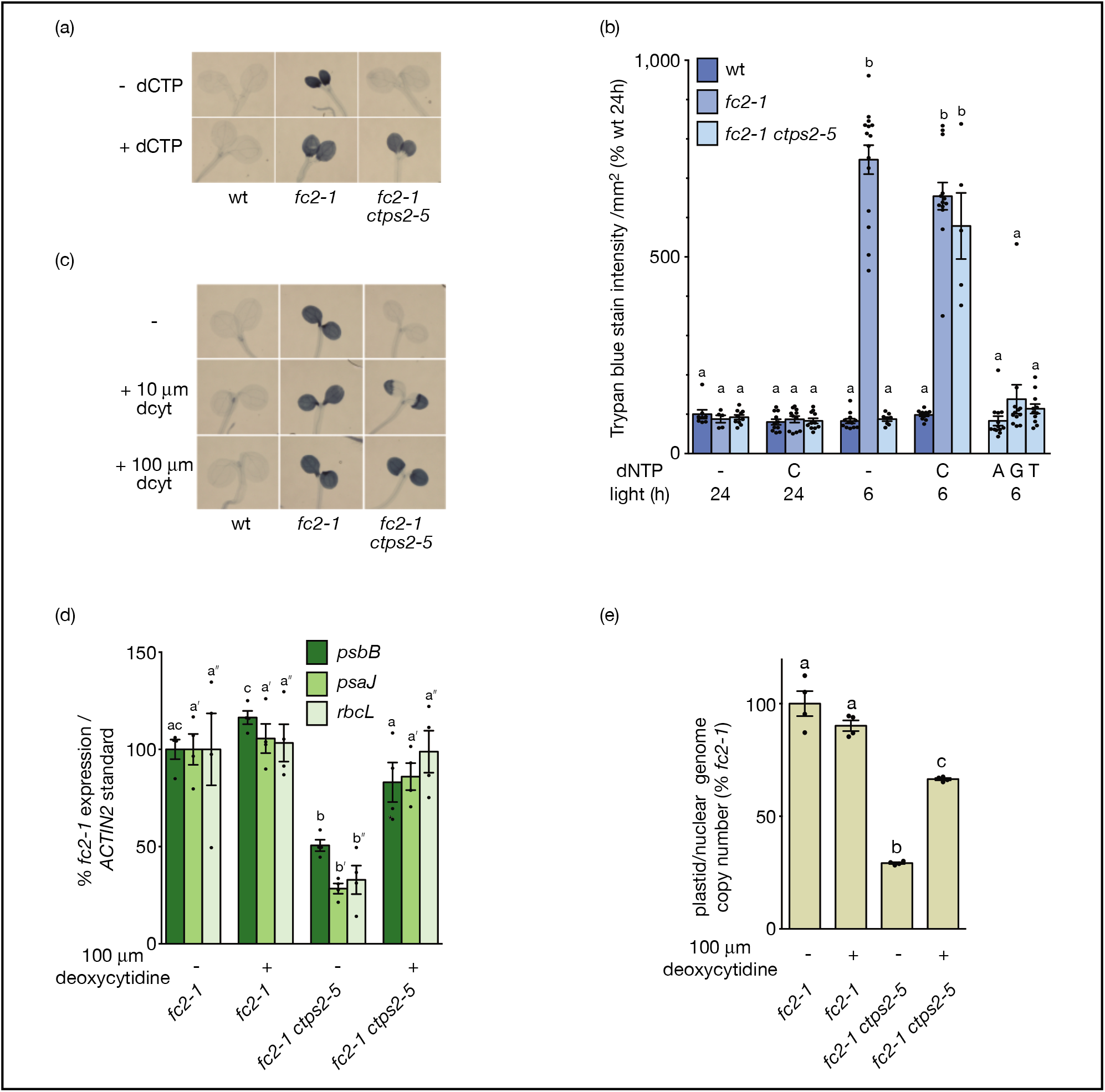
The *ctps2-5* phenotype can be rescued by exogenous dCTP or deoxycytidine feeding. *ctps2-5* mutants were tested for their response to nucleotide and nucleoside feeding. **A)** Shown are representative images of six day-old trypan blue-stained seedlings grown in 6h light / 18h dark diurnal light cycling conditions. The dark blue color is indicative of cell death. Media was supplemented with 2 mM dCTP where indicated. **B)** The mean intensity of the trypan blue staining in A and other seedlings grown in constant 24h light or supplemented with 2 mM of other dNTPs (A = dATP, C = dCTP, G = dGTP, and T = dTTP) (n ≥ 5 biological replicates) +/- SE. **C)** Shown are representative images of six day-old trypan blue-stained seedlings grown in 6h light / 18h dark diurnal light cycling conditions and supplemented with the indicated amounts of deoxycytidine (dcyt). Shown are means of expression levels of **D)** plastid transcripts **E)** and relative plastid genome copy numbers from four day-old seedlings grown in constant light as measured by RT-qPCR (n = 4 biological replicates) +/- SE. Plastid genome copy number was determined by analyzing genomic DNA levels by RT-qPCR using marker probe sets for plastid and nuclear genomes (three sets each). Statistical analyses were performed using one-way ANOVA test. Different letters above bars indicate significant differences within data sets determined by a Tukey-Kramer post-test (p-value ≤ 0.05). In panel D, significance differences for each gene was analyzed separately. ʹ and ʺ designate tests for *psaJ* and *rbcL*, respectively. In all graphs, closed circles represent individual data points.

Next, we tested the ability of the nucleosides cytidine or deoxycytidine to rescue *ctps2-5* phenotypes. Independent of CTPS activity, *Arabidopsis* can synthesize CTP and dCTP from cytidine and deoxycytidine, respectively, through the pyrimidine salvaging pathway (Witte & Herde, 2020). The addition of 100 uM deoxycytidine (but not cytidine) fully restored cell death to *fc2-1 ctps2-5* under light cycling conditions (Figs. 6c, S4). Additionally, deoxycytidine fully restored expression of the plastid transcripts *psbB*, *psaJ*, and *rbcL* (Fig. 6d) and significantly increased plastid DNA copy number in the *fc2-1 ctps2-5* mutant (Fig. 6e). Together, these results suggest the *ctps2-5* mutation suppresses the cell death phenotype of *fc2-1* by reducing a pool of deoxycytidine and/or dCTP specifically available to chloroplasts for DNA synthesis (and indirectly, RNA synthesis).

### *CTPS1-5* have divergent functions in *Arabidopsis*

As *CTPS2* is one of five *CTPS* genes annotated in the *Arabidopsis* genome, we took a phylogenetically informed approach to assess if any of these genes have overlapping roles in chloroplast quality control. We first inferred a phylogeny for CTPS homologs identified in select Brassicaceae, including *Arabidopsis lyrata*, *Capsella rubella*, and *Brassica rapa*, as well as *Tarenaya hassleriana* (a representative of the sister family Cleomaceae), *Gossypium raimondii* (Malvaceae), and *Amborella trichopoda* (a representative of the earliest diverging lineage within the angiosperm lineage). Our tree strongly supports the hypothesis that CTPS1, CTPS4, and CTPS5 share an evolutionary origin that coincides with the origin of the Brassicaceae (Fig. 7a). In addition, the gene that gave rise to CTPS1/4/5 likely descended from a common ancestor that shared a duplication event with CTPS2. CTPS3, however, is monophyletic across angiosperms and the most distantly related CTPS. Thus, if these genes have overlapping roles with CTPS2, CTPS1/4/5 would be the most likely candidates.

**Figure 7.**
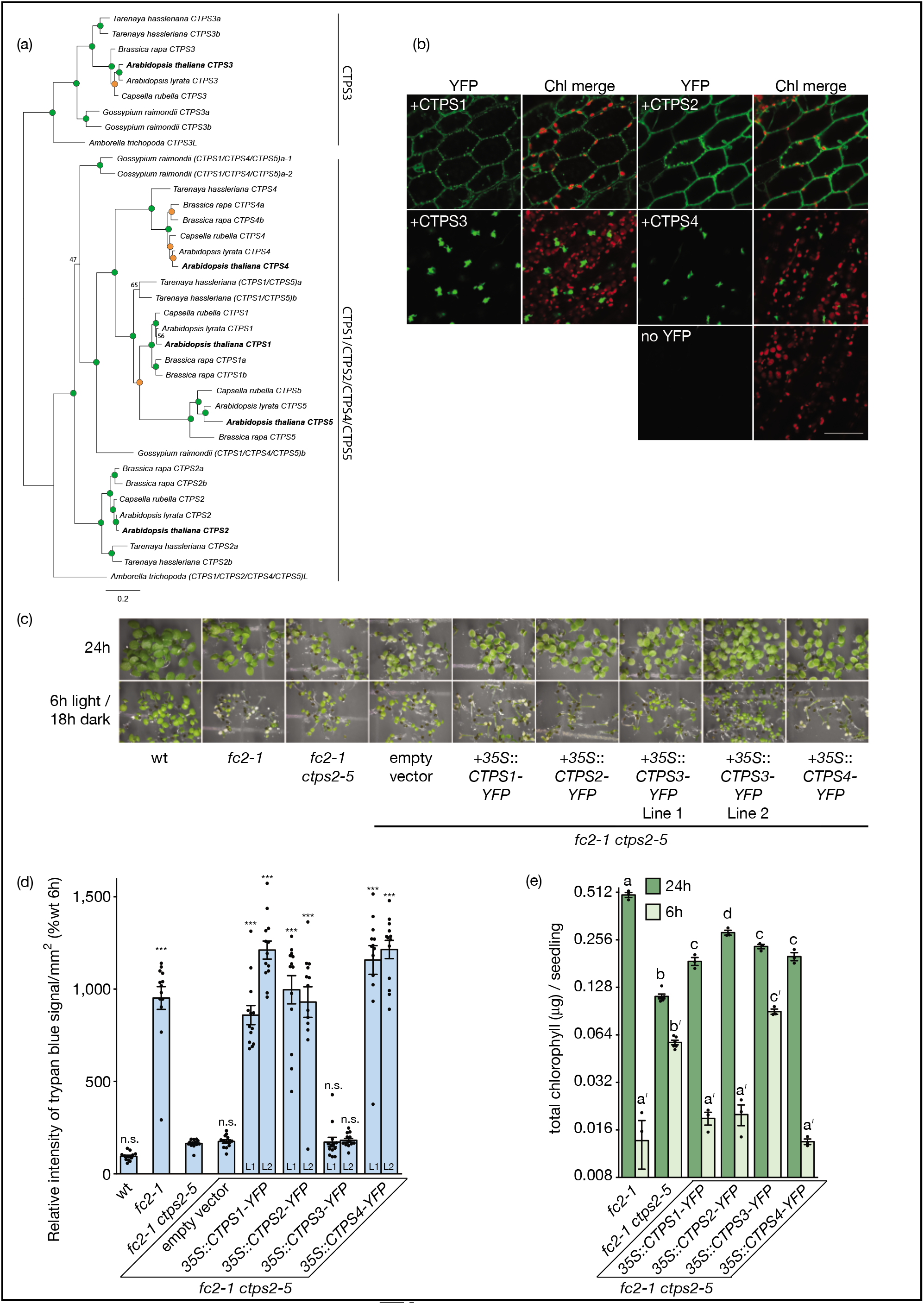
*Arabidopsis* encodes five conserved CTPS enzymes with different functions in chloroplast quality control. An examination of the five CTPS orthologs in *Arabidopsis*. **A)** Maximum likelihood phylogeny of CTPS proteins from select Angiosperms. The monophyletic CTPS3 clade was used to root the tree. Bootstrap scores greater than 95 are shown with green circles, whereas scores between 70-95 are shown with orange. Scores less than 70 are shown. Putative orthologs are labeled based on their relationship to *Arabidopsis CTPS* genes. **B)** Shown are representative images of the localization and morphology of stably overexpressed YFP-tagged CTPS enzymes in four day-old *fc2-1 ctps2-5 Arabidopsis* seedlings grown in constant light. All images acquired using laser scanning confocal microscopy. Scale bar = 40 μm. **C)** Overexpression of *CTPS1-4* was tested for the ability to complement the *ctps2-5* mutant. Shown are pictures of six day-old seedlings grown in the indicated conditions and **D)** means of trypan blue intensity from stained seedlings grown in 6h light / 18h dark diurnal cycling conditions (n = 13 biological replicates) +/- SE. For each *CTPS* overexpression construct, two independent lines were tested and denoted by separate bars (left and right for lines 1 and 2, respectively). Statistical analysis was performed by a one-way ANOVA test followed by Dunnett’s multiple comparisons test with the *fc2-1* ctps2-5 sample. n.s. and *** indicate an adjusted p-value of ≥ 0.05 and ≤ 0.001, respectively. **E)** Chlorophyll content of seedlings in C. Shown are means of biological replicates (n ≥ 3) +/- SE. Statistical analyses for E were performed using one-way ANOVA tests and different letters above bars indicate significant differences within data sets determined by Tukey-Kramer post-tests (p-value ≤ 0.05). Separate statistical analyses were performed for the different light treatments and the significance for the 6h light / 18h dark diurnal cycling values is denoted by letters with a ʹ. In all bar graphs, closed circles represent individual data points.

To test this hypothesis, we first performed a meta-analysis of *CTPS* expression across similar tissues in *Arabidopsis*, *B. rapa*, and *A. trichopoda* using publicly available RNA-seq data (Fig. S5). *Arabidopsis CTPS2* has the highest expression in embryos, consistent with the embryo-lethal phenotype of null mutants (Table S5). *CTPS4* and *CTPS5* are also both highly expressed in ovule tissue, whereas *CTPS1* is most highly expressed in flowers and roots, and *CTPS3* is most highly expressed in leaves. *CTPS* expression is conserved between orthologous loci in *B. rapa*, with some divergence seen in *B. rapa* specific paralogs. Expression of the two *CTPS* genes in *Amborella* is predominantly restricted to reproductive tissues, suggesting a conserved role for *CTPS* genes early in development.

To further examine these proteins, we visualized the subcellular localization of these CTPS enzymes by creating stable *fc2-1 ctps2-5* lines expressing C-terminal YFP-HA-tagged proteins (Fig. 7b). Both CTPS1 and CTPS2 localized throughout the cytoplasm and formed small granule-like aggregates. In strong contrast, CTPS3 and CTPS4 formed large filament-like structures within the cytoplasm and were often near DAPI-stained nuclei (Fig. S6). These large protein filaments were inconsistent in their shapes and sizes and formed both globular and shard-like structures. We were unable to localize CTPS5 as we never recovered any transformants.

To determine which CTPS enzymes may play a role in chloroplast quality control, we tested if these constructs could complement the *ctps2-5* suppressor phenotype (Table S4). As expected, overproduction of CTPS2-YFP-HA restored cell death to the *fc2-1 ctps2-5* mutant under light cycling conditions (Fig. 7c). Overexpression of either *CTPS1* or *CTPS4* also restored cell death suggesting a degree of functional overlap between these three genes. On the other hand, overexpression of *CTPS3* did not restore cell death (17 lines tested). We confirmed these visual phenotypes using trypan blue stains for cell death (Figs. 7d, S7a) and assessing total chlorophyll levels (Fig. 7e). Notably, all four *CTPS* constructs at least partially complemented the pale phenotype of the *fc2-1 ctps2-5* mutant under constant light conditions, suggesting the enzymes were functional (Figs. S7c and e). An immunoblot and RT-qPCR analysis confirmed these lines were producing similarly high levels of the appropriate CTPS-YFP-HA protein and transcripts, respectively (Fig. S7b and c).

To rule out the possibility our *CTPS3* construct was unable to complement due to the particular clone (genomic fragment), promoter (35S), or epitope tag (YFP-HA) we also attempted to complement the *fc2-1 ctps2-5* mutant with cDNA (*CTPS1*-*4*) and genomic (*CTPS5*) clones driven by the *UBQ10* promoter (Daumann *et al.*, 2018) or with a tagless clone of *CTPS3* driven by the *35S* promoter. Complementation with the *UBQ10* promoter constructs led to very similar results; *CTPS1*-, *2*-, and *4*-*YFP* complemented the cell death phenotype under light cycling conditions, while the *CTPS3-YFP* clone did not (Fig. S7d). In this case, we were able to isolate lines expressing *CTPS5-YFP*, which led to the complementation of the *ctps2-5* phenotype. An immunoblot demonstrated these lines accumulated similar levels of CTPS-YFP protein (Fig. S7e). Finally, overexpressing untagged CTPS3 protein (*35S::CTPS3* construct) failed to restore cell death to *fc2-1 ctps2-5* (25 lines tested), confirming the inability of *CTPS3* to replace *CTPS2* function (Fig. S7f). We confirmed overexpression of *CTPS3* in these lines by RT-qPCR (Fig. S7g). Together, these results support our evolutionary analysis that CTPS1/2/4/5 and CTPS3 represent two groups of enzymes with conserved and distinct functions.

### CTPS3 reveals a link between chloroplast quality control and plastid DNA content

Next, we wanted to understand why CTPS3 was unable to compensate for CTPS2 function. One possible reason is CTPS3 may have a distinct subcellular localization preventing nucleotide allocation to chloroplasts. Although the filamentous localization of CTPS3 is clearly distinct from the evenly distributed cytoplasmic localization of CTPS2, it is similar to CTPS4, which can compensate for the lack of CTPS2 (Fig. 7c). To test if CTPS3 and CTPS4 are co-localizing to the same filament-like structures, we created lines producing CFP-tagged CTPS3 or CTPS4 proteins, crossed them to lines producing YFP-tagged CTPS3 or CTPS4 proteins, and imaged the resulting F1 generation. When co-expressed, CTPS3-YFP and CTPS4-CFP (or with reversed tags) clearly co-localized to the same cytoplasmic filaments in cells (Fig. 8a) demonstrated by the overlap of YFP and CFP signal. Interestingly, the shape of these structures appears to depend on the type of tag and if proteins are produced alone or together (Fig. S8a). When individually expressed with CFP tags, the proteins produce shard-like structures that are mostly absent in lines expressing the YFP-tagged forms. When CTPS3 and CTPS4 are co-expressed, they localize to much smaller circular puncta, suggesting these filaments are variable and dependent on CTPS protein levels. Overall, however, the localization of these proteins does not account for their different functions in the cell.

**Figure 8.**
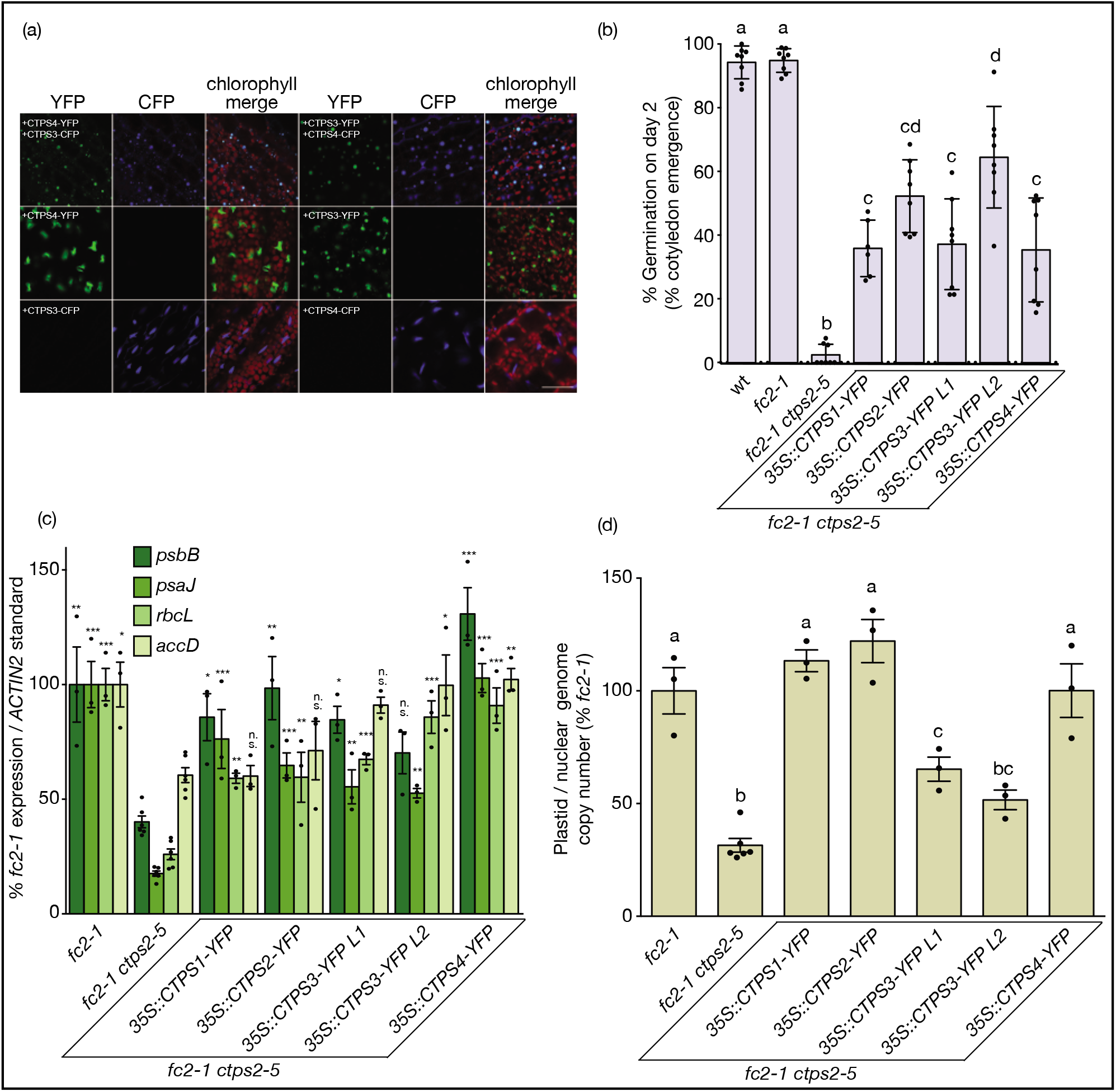
*CTPS1/2/3/4* have different effects on plastid DNA copy number. *Arabidopsis* CTPS orthologs were tested for co-localization, effect on germination, and contribution to plastid gene expression and genome maintenance. **A)** Shown are representative images of the localization and morphology of stably overexpressed YFP- and CFP-tagged CTPS enzymes in four day-old *Arabidopsis* seedlings grown in constant light. All images acquired using laser scanning confocal microscopy. Scale bar = 40 μm. **B)** % germination of seedlings two days post stratification. Shown are the average germination rates of various genotypes per set of 50-100 seeds (n = 4 sets) +/- SE. Germination was defined as cotyledon emergence from the seed coat. **C)** Shown are means of expression levels of plastid transcripts from four day-old seedlings grown in constant light as measured by RT-qPCR (n ≥ 3 biological replicates) +/- SE. **D)** Shown are the means of relative plastid genome copy number from four day-old seedlings grown in constant light (n ≥ 3 biological replicates) +/- SE. Values were determined by analyzing genomic DNA levels by qPCR using marker probe sets for plastid and nuclear genomes (three sets each). Statistical analyses for B and D were performed by a one-way ANOVA test and different letters above bars indicate significant differences determined by a Tukey-Kramer post-test (p-value ≤ 0.05). Statistical analysis for C was performed by a one-way ANOVA test followed by Dunnett’s multiple comparisons test with the *fc2-1* sample. *, **, ***, and n.s. indicate an adjusted p-value of ≤ 0.05, ≤ 0.01, ≤ 0.001, and ≥ 0.05, respectively. In all bar graphs, closed circles represent individual data points.

Next, we tested the ability of *CTPS* overexpression constructs to complement the delayed germination phenotype of the *fc2-1 ctp2-5* mutant. In this case, overexpression of *CTPS1*-*4* all partially rescued this delayed germination (Figs. 8b, S8b). This suggests the delayed germination phenotype of *ctps2-5* is due to a general decrease in CTP synthesis and is not linked to chloroplast quality control or any particular CTPS enzyme.

Finally, we tested the ability of the *CTPS* overexpression constructs to rescue the reduced plastid gene expression and DNA copy number in *fc2-1 ctps2-5* mutants. Interestingly, *CTPS1-4* all partially restored plastid gene expression (at least three of the four genes monitored were significantly increased by each construct compared to *fc2-1 ctps2-5*) (Fig. 8c). However, only *CTPS1*, *2*, and *4* restored plastid DNA copy number to *fc2-1* levels (Fig. 8d). Overexpression of *CTPS3* did not. All together, these results suggest that the conserved CTPS1/2/4/5 clade can provide adequate dCTP levels for chloroplast DNA synthesis required for chloroplast quality control signals, while CTPS3 cannot.

## Discussion

During photosynthesis, chloroplasts experience stress in the form of ROS that damages molecules within these essential compartments. These ROS can act as signals to regulate chloroplast turnover, cell death, acclimation responses, and retrograde signaling to the nucleus (Chan *et al.*, 2015; Kleine & Leister, 2016; de Souza *et al.*, 2017; Rochaix & Ramundo, 2018; Dogra & Kim, 2019). Although decades of genetic research with *Arabidopsis* have demonstrated multiple chloroplast signaling pathways exist, the molecular mechanisms behind them have mostly remained obscure.

Here we report the characterization of the *fc2-1* suppressor *ctps2-5*, which affects CTPS2, an essential enzyme involved in cytoplasmic CTP synthesis. In this mutant, ^1^O_2_ accumulation is uncoupled from chloroplast degradation, cell death, and retrograde signaling (Figs. 1a, 2b, 3b, and 4B) suggesting the ^1^O_2_ signal has been blocked. In addition, plastid gene expression and genome copy number are reduced, leading to delayed chloroplast development. While lowered plastid gene expression could be attributed to reduced plastid CTP pools for RNA synthesis, our feeding experiments indicated *ctps2-5* plastids are limited for dCTP (which can be indirectly synthesized from CTP) and DNA synthesis. Feeding of dCTP or deoxycytidine (which can be converted to dCTP through the pyrimidine salvage pathway) both restored cell death in *fc2 ctps-5* mutants while deoxycytidine was able to restore plastid gene expression and plastid DNA copy number. Recovery of cell death with deoxycytidine is mostly likely due to restoring dCTP rather than CTP synthesis, as CTP cannot be synthesized from deoxycytidine without CTPS activity (Witte & Herde, 2020). Furthermore, direct feeding of cytidine was unable to restore cell death to *fc2-1 ctps2-5* mutants. As low plastid DNA levels can limit plastid gene expression (Sasaki *et al.*, 1990; Udy *et al.*, 2012), we hypothesize reduced dCTP availability is at least partly responsible for the reduced plastid transcripts. Finally, we observed a block of cell death in *fc2-1* seedlings when they were grown in the presence of two different DNA gyrase inhibitors (Figs. 5c and d), suggesting that lowered plastid DNA content is sufficient to suppress *fc2-1* phenotypes. Therefore, we conclude that *ctps2-5* suppresses ^1^O_2_ signaling by reducing plastid DNA copy number.

Previously, we demonstrated specific mutations that reduce PEP-dependent gene expression can block the ^1^O_2_ signal (e.g., *ppr30* and *mTEF9*), possibly by affecting transcripts encoding signaling factors (Alamdari *et al.*, 2020). By reducing both plastid DNA content and transcript levels, the *ctps2-5* mutation may block chloroplast degradation through a similar mechanism. Moreover, like *ppr30* and *mteff9* mutants, *ctps2-5* single mutants also suffer less cell death under EL stress further suggesting that they all affect a common pathway activated under naturally induced photo-oxidative stress (Fig. 4d). Such a pathway appears genetically distinct from the EXECUTOR (EX) pathway (Wagner *et al.*, 2004) that mediates some chloroplast ^1^O_2_ signaling to the nucleus; *ex1* mutants are unable to suppress *fc2-1* phenotypes (Woodson *et al.*, 2015). As such, this work suggests chloroplast quality control signals can be regulated within the cytoplasm through the synthesis or salvaging of nucleotide pools. This may permit cells to control the fates of their organelles, particularly during photomorphogenesis when the RNA and DNA content of plastids is rapidly increasing.

Strikingly, although plastid function is impaired in the *ctps2-5* mutant, mitochondrial gene expression (Fig. S3) and genome copy number (Fig. 5b) appear unaffected. This suggests CTPS2 provides a specific pool of nucleotides dedicated to chloroplast DNA synthesis and aligns with previous reports hinting at the existence of organelle-specific nucleotide pools. Mutations that affect *VENOSA4/VEN4*, which encodes a nuclear-localized NTPase that cleaves dNTPS to their nucleosides, lead to pale phenotypes and a reduction of photosynthetic protein content (Xu *et al.*, 2020). These phenotypes are rescued by the feeding of dCTP (but not other dNTPs) suggesting chloroplasts may be particularly sensitive to a depletion of deoxycytidine-based nucleotides. *cls8* mutants, which are defective in the cytoplasmic enzyme RIBONUCLEOTIDE REDUCTASE 1 (RNR1) that reduces ribonucleotides to their deoxyribonucleotide counterparts, have variegated phenotypes with bleached sectors in the leaves, reduced plastid DNA, impaired plastid DNA replication, and impaired chloroplast division (Garton *et al.*, 2007). Outside of DNA metabolism, CTP and dCTP are also essential cofactors to produce CDP- and dCDP-choline, intermediates in the biosynthesis of phospholipids for chloroplasts and other cellular components. Mutants defective in these pathways have defects in photoautotrophic growth and undeveloped chloroplasts (Haselier *et al.*, 2010; Hong *et al.*, 2018). Therefore, it is possible such dCTP-dependent metabolites may also play a role in the *ctps2-5* phenotype or chloroplast quality control pathways.

How could the CTPS2 enzyme, localized to the cytoplasm, have such a strong effect on chloroplast function while seemingly being dispensable for mitochondrial RNA and DNA metabolism? We sought to answer this question by investigating the other four *Arabidopsis* CTPS orthologs. Our phylogenetic analysis suggested that while all five enzymes are conserved in the

Brassicaceae, CTPS2 is a member of an ancient clade including CTPS1/4/5 that is distinct from CTPS3 (Fig. 7a). The deep coalescence between CTPS3 and the CTPS2/1/4/5 clade suggested these two groups of enzymes have evolved distinct functions in flowering plants. To test this hypothesis directly, we examined the ability of the other CTPS enzymes to complement the *ctps2-5* phenotype. Consistent with the phylogenetic tree, cell death was restored in the *fc2-1 ctps2-5* mutant by overexpression of any member of the CTPS2 clade (*CTPS1/2/4/5*) while overexpression of *CTPS3* did not (Figs 7c and S7d). Furthermore, only *CTPS1/2/4* were able to fully restore plastid DNA copy number (*CTPS5* was not tested). Surprisingly, *CTPS1-4* at least partially restored expression of the four plastid genes tested. It is possible *CTPS3* does not fully restore all plastid gene expression as some transcripts are more sensitive to DNA content than others (Fig. 5a) (Sasaki *et al.*, 1990; Udy *et al.*, 2012). An exciting alternative possibility is lowered plastid DNA content may block ^1^O_2_ signaling through an unknown mechanism. In either case, CTPS3 does not appear to contribute to the pool of CTP used to provide dCTP to chloroplasts. Instead, that function may be fulfilled by the conserved CTPS1/2/4/5 clade of enzymes.

Although consistent with evolving separate functions, the inability of CTPS3 to operate in chloroplast quality control pathways was still unexpected. CTPS3 has been previously shown to be a bonafide CTPS enzyme *in vitro* (Daumann *et al.*, 2018). Here, CTPS3 appears functional *in vivo* as the reduced plastid transcripts and late germination phenotypes of *ctps2-5* were partially rescued by its overproduction (Fig. 7d). Lastly, in *Arabidopsis* and *B. rapa*, *CTPS3* is predominately expressed in leaves, indicating a potential role in chloroplast function in these species. However, the dominant tissue of expression for *CTPS3* in *Amborella* is in reproductive tissues, suggesting its ancestral function may be independent of chloroplast biogenesis or photosynthesis

How a CTPS enzyme could allocate nucleotides to a cellular sub-compartment is unknown. One possibility is their different cytoplasmic localizations could drive the synthesis of distinct metabolite pools. As shown here (Fig. 7b) and previously (Daumann *et al.*, 2018), CTPS1 and 2 localize throughout the cytoplasmic compartment. CTPS3-5, on the other hand, form large filament-like structures in the cytoplasm, with CTPS3 and CTPS4 co-localizing to the same foci (Fig. 8a). Observed in bacteria, fungi, *Drosophila*, and humans, the significance behind these filament structures (dubbed “cytoophidia”) is still not understood (Liu, 2016). Although the function of these filaments may vary between species (e.g., to regulate activity), there is no correlation with chloroplast quality control. The mechanism behind CTP allocation, therefore, cannot be completely explained by localization or filament formation.

The duplication and retention of *CTPS* genes encoding an essential protein for primary nucleotide metabolism suggests CTP synthesis in plants is complex. In addition to different functions and sub-cellular localizations, their expression patterns vary greatly (Fig. S5) and are induced by different types of stress: *CTPS1* by salt stress (Zimmermann *et al.*, 2004), *CTPS2* during cold acclimation (Oono *et al.*, 2006) and cadmium stress (Sarry *et al.*, 2006), and *CTPS4* by drought (Zimmermann *et al.*, 2004). Thus, by having five orthologs, plants may use a mechanism to provide dCTP (and possibly CTP) to different compartments in a cell, each with its own fluctuating demand for nucleotides. A deeper understanding of these enzymes’ functions may help us to understand how cells regulate nucleotide allocation with chloroplast stress and quality control pathways.

## Conclusions

As centers of energy production, chloroplasts are essential for plant life, and by extension, for most life on the planet. Assembling chloroplasts and their photosynthetic machinery is an extremely complex process involving a suite of proteins, whose levels and functions are controlled by chloroplast signaling pathways. Because chloroplasts experience inevitable photo-oxidative damage due to photosynthesis’s unique chemistry, these organelles must have chloroplast quality control mechanisms. These signals not only repair individual chloroplasts, but remove severely damaged chloroplasts to protect the cell from further oxidative damage. Here we show modulation of cytoplasmic nucleotide synthesis by a conserved clade of enzymes is a mechanism by which cells can control chloroplast stress signaling through shifts in plastid DNA content and gene expression. This not only provides further insight to chloroplast development and stress signaling, but how metabolite pools in the cytoplasm may be diverted to specific subcellular components for growth and development. A deeper understanding of these cellular mechanisms will be essential for developing tools to increase crop production and yield under stressful and dynamic environments.

## Supporting information

Alamdari et al supporting information

Alamdari et al Table S3

## Acknowledgements

We thank Dr. Hans-Henning Kunz (Washington State University) for generously sharing the *UBQ10::CTPS* vector constructs, Dr. Ramin Yadegari (University of Arizona) for providing a fluorescent microscope for phenotyping transgenic lines, Dr. David Baltrus (U. Arizona) for providing a plate reader for chlorophyll measurements, and Tania Chakraborty (University of Arizona) for proving technical assistance with RNA extraction, cDNA synthesis, and PCR. The authors acknowledge the Division of Chemical Sciences, Geosciences, and Biosciences, Office of Basic Energy Sciences of the U.S. Department of Energy grant DE-SC0019573 and UA Core Facilities Pilot Program grant awarded to J.D.W, NSF-IOS grant 1758532/2021753 awarded to A.D.L.N., and the NSF Graduate Research Fellowship Grant DGE-1746060 awarded to K.R.P. The authors have no conflict of interest to declare.

## Author Contributions

KA performed all cloning, RT-qPCR, physiological growth experiments, and plant treatments. KEF performed all microscopy experiments and their sample preparations. JDW performed the SOSG assays, Pchlide measurements, and cell death assays. DWW developed and performed chlorophyll assays. SR performed all protein extractions and immunoblots. KRP and ADLN performed phylogenetic and in silico expression analyses. KA, KEF, DWW, ADLN, and JDW conceived all experiments and analyzed the data. JDW conceived original scope of project, managed the project, and wrote the manuscript. All authors contributed in reviewing the manuscript and approved of final version.

### Short legends for supporting information

Methods

S1. Bacterial growth conditions

S2. Construction of complementation vectors

S3. Protein Extraction and Immunoblotting

Supplemental Tables

Table S1. Primers use in this study

Table S2. Vectors used in this study

Table S3. List of mutations in the mapped region of the *fc2-1 fts39* line

Table S4. Complementation and phenocopy analyses of overexpression vectors

Table S5. *ctps2* alleles

Supplemental Figures

Figure S1. A loss of function *ctps2* allele suppresses cell death in the *fc2-1* mutant

Figure S2. Effect of the *ctps2-5* mutation on chloroplast compartment size

Figure S3. The *ctps2-5* mutation does not significantly affect mitochondrial gene expression

Figure S4. The *ctps2-5 fts* phenotype can be rescued by exogenous deoxycytidine feeding Figure S5. *CTPS1-5* have distinct expression patterns across flowering plants

Figure S6. *Arabidopsis* CTPS3 and CTPS4 are extra-nuclear localized.

Figure S7. The five *Arabidopsis* CTPS enzymes have different functions in chloroplast quality control

Figure S8. Effect of *CTPS* overexpression on CTPS protein localization and germination

### Data Statement

All data used in the manuscript are included within. All biological materials generated for this work will be available upon request.

