## Supplementary material for "Chloroplast quality control pathways are dependent on plastid DNA synthesis and nucleotides provided by cytidine triphosphate synthase two": Alamdari et al supporting information

### Methods

#### S1. Bacterial growth conditions

All bacteria (*E. coli* and *Agrobacterium tumefaciens* strains) were grown in liquid Miller nutrient broth or solid medium containing 1.5% agar (w/v). Cells were grown at 37°C (*E. coli*) or 28°C (*A. tumefaciens*) with the appropriate antibiotics and liquid medium was shaken at 225 rpm.

#### S2. Construction of complementation vectors

DNA fragments were amplified from wt *Arabidopsis* genomic DNA template using the Q5 High Fidelity DNA Polymerase (New England Biolabs). These DNA fragments were gel-purified using the Zymoclean Gel DNA Recovery Kit (Zymo Research) and cloned into the Gateway compatible vector pENTR-D/TOPO or pDONR221 (Invitrogen) according to the manufacturer's instructions. All primers and vectors are listed in Tables S1 and S2, respectively.

Using LR clonase (Invitrogen) according to manufacturer's instructions, DNA fragments were then transferred to either pEARLEYGATE101 (35S overexpression promoter plus C-terminal HA and YFP tags (Earley *et al.*, 2006)) or pEARLEYGATE102 (35S overexpression promoter plus C-terminal HA and CFP tags (Earley *et al.*, 2006)) to produce fluorescently tagged protein driven by the constitutive 35S promoter. To express native CTPS2 protein with the *CTPS2* promoter, the *CTPS2* genomic fragment was transferred to pGBGWY (Zhong *et al.*, 2008). The completed vectors were then transformed into the *A. tumefaciens* strain GV301 (with the pSOUP helper plasmid for vectors with the pGBGWY backbones). These strains were then subsequently used to transform *Arabidopsis* via the floral dip method. $T_1$ plants were selected for their ability to grow on Basta-soaked soil and propagated to the next generation. $T_2$ lines were monitored for single insertions (segregating 3:1 for Basta resistance:sensitivity) and propagated to the next generation. Finally, homozygous lines were selected in the $T_3$ generation based on 100% Basta-resistance. In the case of lines expressing YFP- or CFP-tagged constructs, lines were selected that exhibited strong and evenly distributed YFP or CFP signal (as observed under a fluorescent microscope). The *fts* phenotypes of all transgenic lines were scored and the results are listed in Table S4. Representative lines were further characterized and included in the manuscript.

### S3. Protein Extraction and Immunoblotting

Seven day-old seedlings were frozen in liquid $N_2$ and homogenized in extraction buffer containing 50 mM Tris-HCl (pH 7.5), 150 mM NaCl, 0.5% IGEPAL (v/v), and Pierce protease inhibitor (Thermo Scientific). The mixture was vortexed and incubated on ice for 30 min. Cell debris was twice removed by centrifugation (8,000 x g for 10min at 4ºC). Total protein concentration was measured using a Bradford assay (Bio-Rad). 15 ug protein from each sample was separated on a 4-20% Tris-Glycine gel (Mini-PROTEAN TGX gel, Biorad) and electrotransferred to PVDF membrane (Millipore Sigma) at 100 V for 60 min. The blots were probed with anti-GFP (Invitrogen) and anti-Actin (Sigma) primary antibodies. Membranes were then incubated with horseradish peroxidase conjugated secondary antibodies (Biorad). Detection was performed using the ProSignal Pico ECL kit (Prometheus). Gels were imaged on a ChemiDoc XRS+ System with Image Lab Software (BioRad).

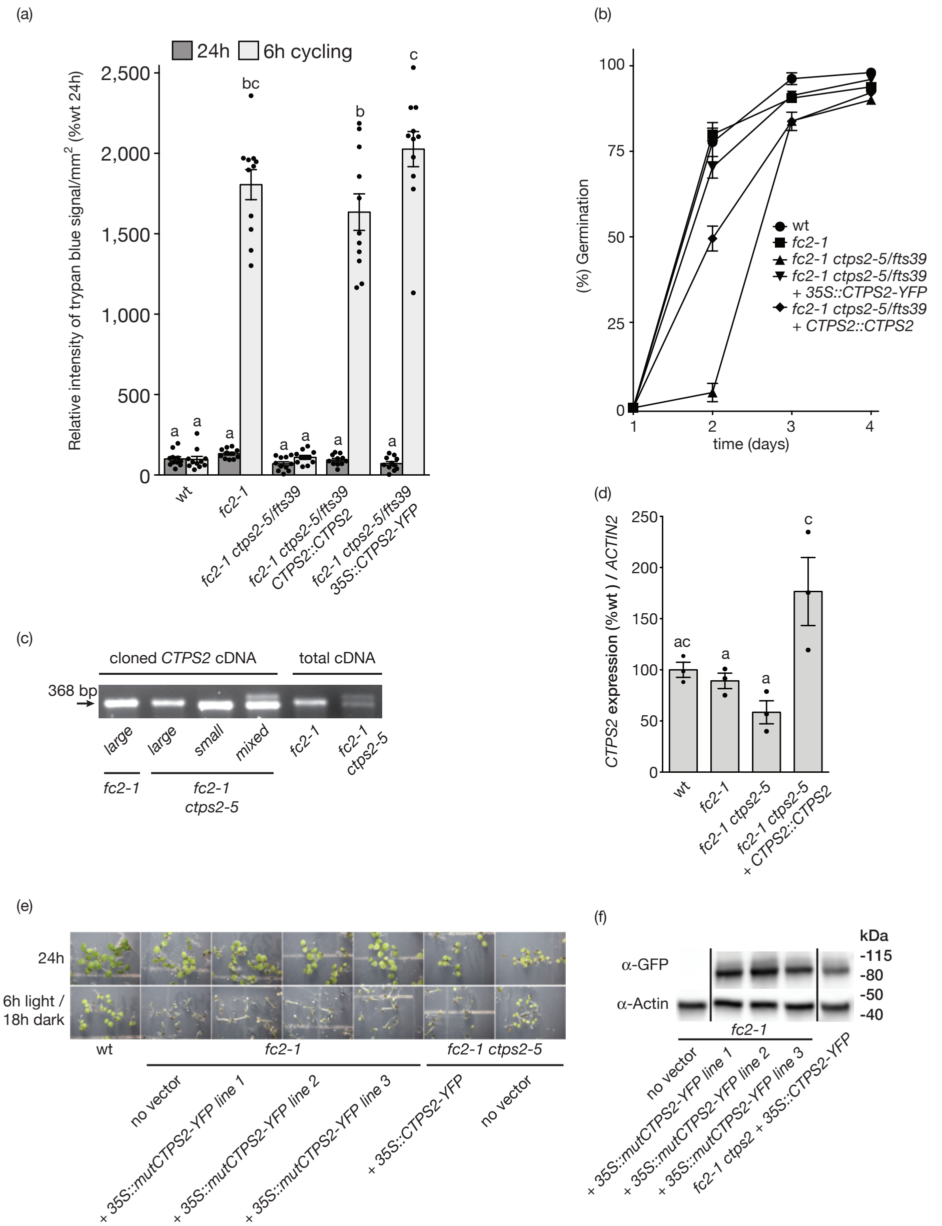


**<u>Fig. S1 legend. A loss of function *ctps2* allele suppresses cell death in the *fc2-1* mutant.</u>**

**A)** Quantification of trypan blue staining (relative intensity/mm$^2$) of six day-old seedlings grown in constant (24h) light or 6h light / 18h dark diurnal (6h) cycling conditions from Fig. 1B. Shown are means of biological replicates (n = 11) +/- SE. **B) %** germination of seedlings (days post stratification). Shown is the average % germination of various genotypes per set of 50-100 seeds (n = 4 sets) +/- SE. Germination was defined as cotyledon emergence from the seed coat. **C)** Shown are PCR amplified DNA products from cloned cDNA fragments or total cDNA. In lane four, vectors containing both small and large cDNA fragments were mixed (1:1) prior to PCR amplification. **D)** Shown are means of expression levels of *CTPS2* transcripts from four day-old seedlings grown in constant light as measured by RT-qPCR (n = 3 biological replicates) +/- SE. **E)** Shown are five day-old seedlings grown under constant (24h) light or 6h light / 18h dark diurnal conditions. **F)** An immunoblot of proteins extracted from the same seedlings in E. CTPS-YFP and actin levels were determined using anti-GFP and anti-actin antibodies, respectively. Statistical analyses were performed by one-way ANOVA tests and different letters above bars indicate significant differences determined by a Tukey-Kramer post-test ($p$-value $\leq 0.05$). In all bar graphs, closed circles represent individual data points.

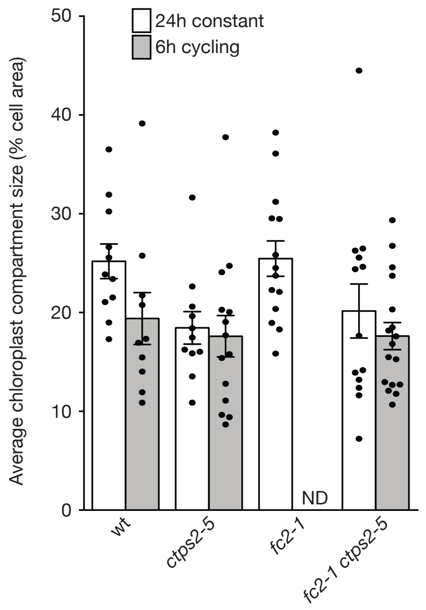


**Fig S2 legend. Effect of the *ctps2-5* mutation on chloroplast compartment size.**
Shown are means of the average % chloroplast compartment size (total chloroplast area/cell area) ($n \geq 10$ cells) of four day-old seedlings grown in constant (24h) light or 6h light / 18h dark diurnal cycling (6h) conditions as assessed by TEM (+/- SE). Chloroplast volume in *fc2-1* seedlings grown in 6h light conditions was not determined (ND) due to extensive cellular degradation. A Statistical analysis were performed by a one-way ANOVA test followed by a Tukey-Kramer post-test. No significant differences were calculated between samples for a single genotype or among genotypes across a given light condition (p-value $\geq 0.50$).

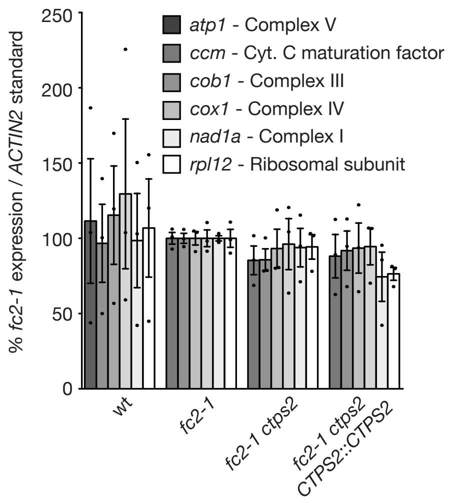


**Fig S3 legend. *The ctps2-5* mutation does not significantly affect mitochondrial gene expression.**

Shown are mean expression levels of mitochondrial transcripts from four day-old seedlings grown in constant light as measured by RT-qPCR (n = 3 biological replicates) +/- SE. This experiment used the same cDNA samples in Fig. 2 that were used to measure plastid transcripts. Closed circles represent individual data points. Statistical analysis was performed by a one-way ANOVA test followed by Dunnett's multiple comparisons test with the *fc2-1* sample. The expression of each transcript was not significantly different between any genetic background (p-value $\geq 0.60$).

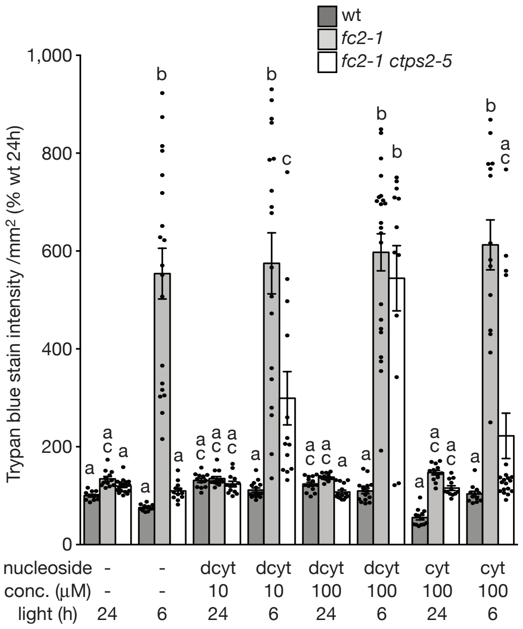


**<u>Fig S4 legend. The *ctps2-5 fts* phenotype can be rescued by exogenous deoxycytidine feeding.</u>**

Shown are the mean values of trypan blue stain intensity ($mm^2$) across the cotyledons of six day-old seedlings (% wt in 24h light) ($n \geq 11$) +/- SE. Seedlings were grown in either constant light (24h) or 6h light / 18h dark diurnal cycling (6h) conditions and supplemented with the indicated amounts of cytidine (cyt) or deoxycytidine (dcyt). Statistical analyses were performed using a one-way ANOVA test. Different letters above bars indicate significant differences within data sets determined by a Tukey-Kramer post-test (p-value $\leq 0.05$). Closed circles represent individual data points.

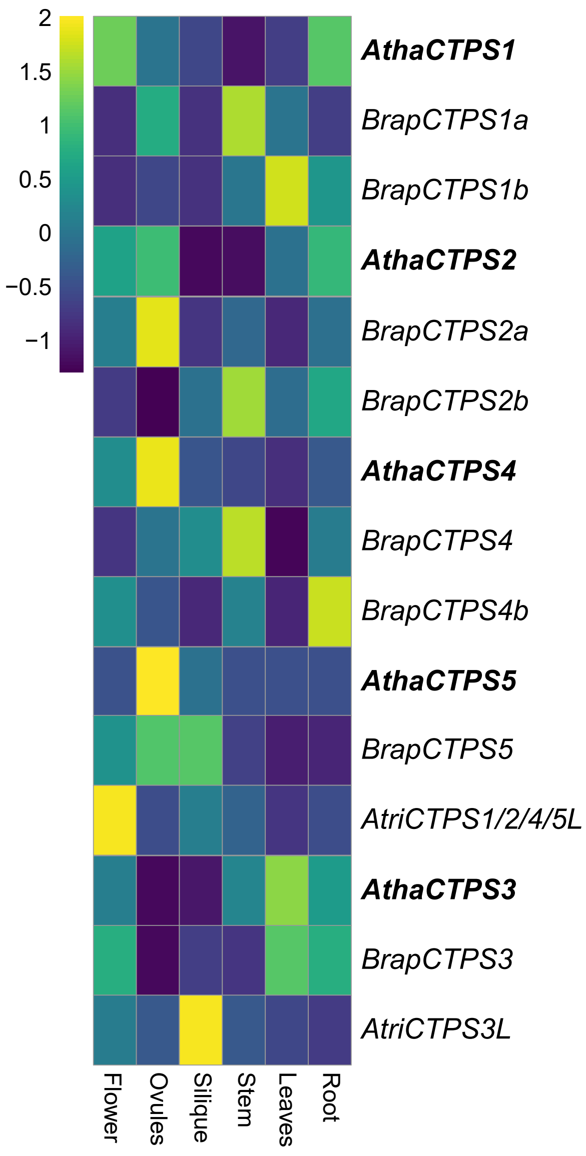


**Fig. S5 legend. *CTPS1-5* have distinct expression patterns across flowering plants.** Z-score normalized expression levels for *Arabidopsis*, *Brassica rapa*, and *Amborella trichopoda CTPS* putative orthologs from six different tissues. *CTPS* nomenclature is adopted from Fig. 7a.

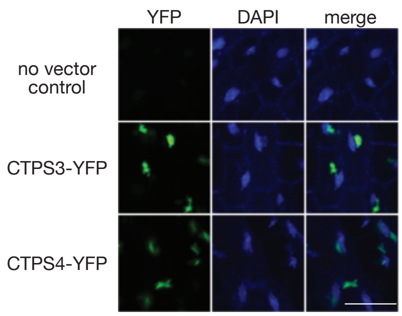


**Fig. S6 legend. *Arabidopsis* CTPS3 and CTPS4 are extra-nuclear localized.**
The subcellular localization of YFP-tagged CTPS protein was assessed in DAPI-stained seedlings. Shown are representative images of CTPS-YFP enzymes (green) and DAPI-stained nuclei (blue) in four day-old *fc2-1 ctps2-5 Arabidopsis* seedlings grown in constant light. All images acquired using laser scanning confocal microscopy. Scale bar = 40 μm.

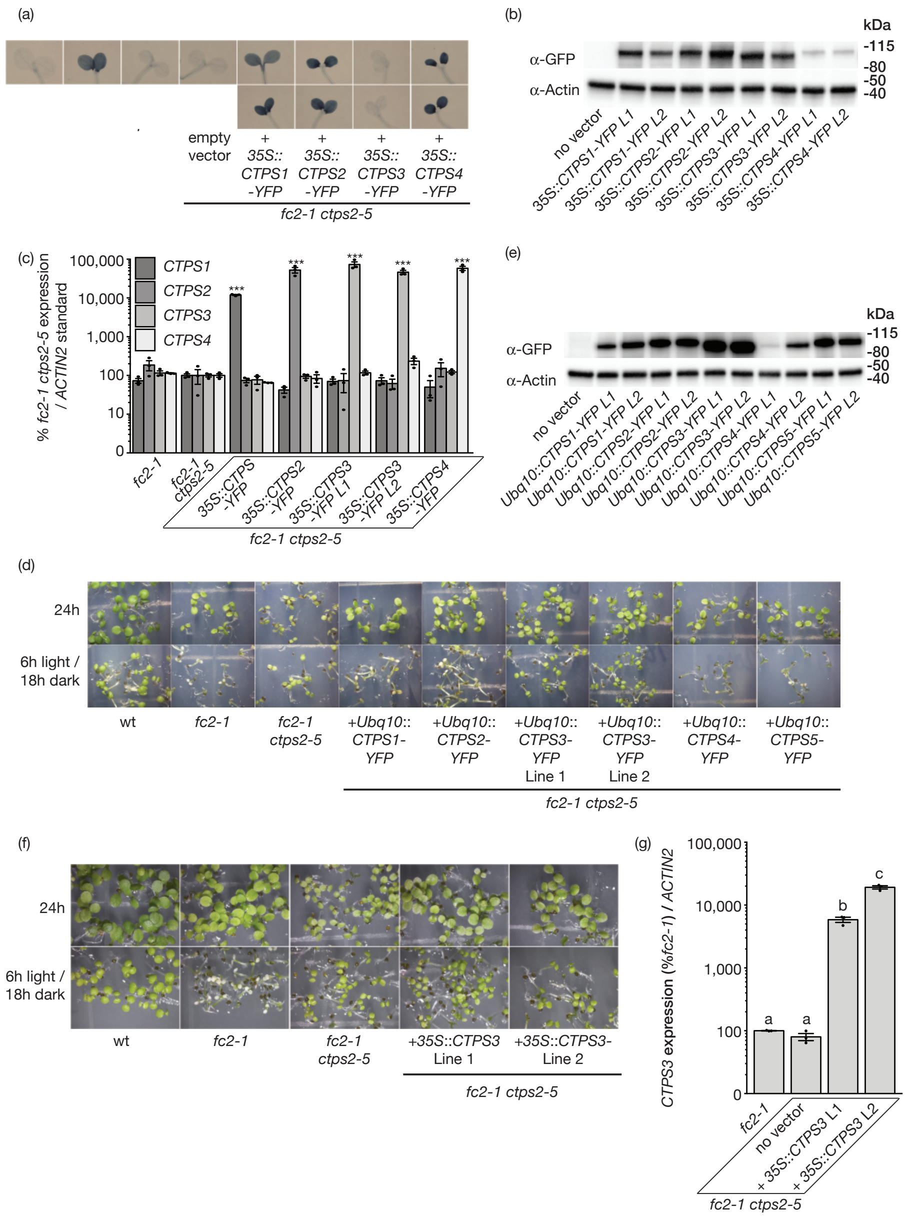


**<u>Fig. S7 legend. The five *Arabidopsis* CTPS enzymes have different functions in chloroplast quality control.</u>**

The five *Arabidopsis CTPS* orthologs were tested for their ability to complement the *ctps2-5* phenotypes in 6h light / 18h dark diurnal cycling conditions. **A)** Cell death was assessed by staining the seedlings shown in Fig. 7C with trypan blue. Shown are pictures of these trypan blue-stained seedlings quantified in Fig. 7D. For each *CTPS* overexpressing construct, two independent lines were tested. The top and bottoms rows are lines 1 and 2, respectively. The deep dark blue color is indicative of cell death. **B)** An immunoblot of proteins extracted from the same seedlings grown in constant light conditions (Fig. 7C). CTPS-YFP and actin levels were determined using anti-GFP and anti-actin antibodies, respectively. **C)** Shown are means of *CTPS1/2/3/4* transcripts in *CTPS* OX lines used in Figs 7 and 8 (line 1 of each and line 2 of *35S*::*CTPS3*) as measured by RT-qPCR +/- SE. RNA was extracted from four day-old seedlings grown in constant light (n = 3 biological replicates). A statistical analysis was performed by a one-way ANOVA test followed by Dunnett's multiple comparisons test with the *fc2-1 ctps2-5* sample. *** indicates an adjusted p-value of $\leq$ 0.001. **D)** Shown are six day-old seedlings grown in constant light (24h) and 6h light / 18h dark diurnal cycling conditions. The seedlings here harbor constructs containing cDNA clones of *CTPS* genes (except for *CTPS5*, which is a genomic fragment) driven by the *UBQ10* promoter. **E)** An immunoblot of proteins extracted from the seedlings in panel D grown in constant light conditions. CTPS-YFP and actin levels were determined using anti-GFP and anti-actin antibodies, respectively. **F)** Shown are six day-old seedlings grown in constant (24h) light and 6h light / 18h dark diurnal cycling conditions. The seedlings here harbor constructs over-producing native (tagless) CTPS3 protein. **G)** Shown are means of expression levels of *CTPS3* transcripts seedlings in panel F grown in constant light as measured by RT-qPCR (n = 3 biological replicates) +/- SE. Statistical analysis was performed by a one-way ANOVA test and different letters above bars indicate significant differences determined by a Tukey-Kramer post-test (p-value $\leq$ 0.05). In all graphs, closed circles represent individual data points.

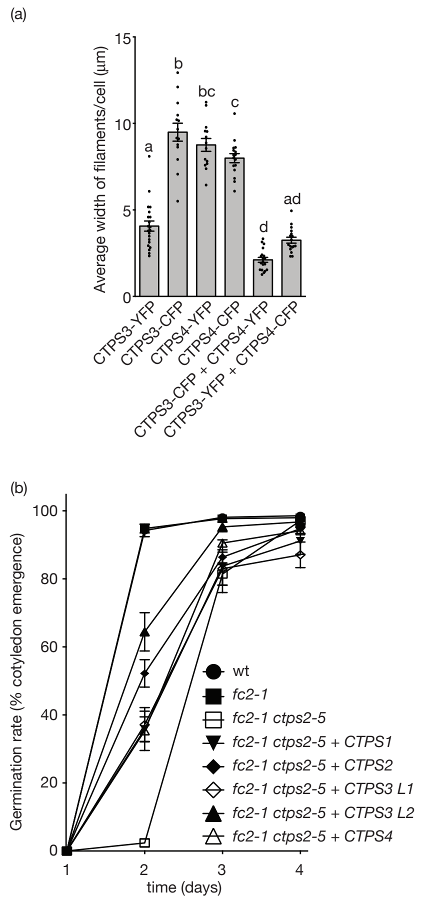


**Fig. S8 legend. Effect of *CTPS* overexpression on CTPS protein localization and germination.**

**A)** Shown are the average maximum widths per cell of the protein filaments visualized in Fig. 8A (n = 11) +/- SE. Statistical analysis was performed by a one-way ANOVA test and different letters above bars indicate significant differences determined by a Tukey-Kramer post-test (p-value $\leq$ 0.05). **B) %** germination of seedlings (days post stratification). Shown are the average germination rates of various genotypes per set of 50-100 seeds (n = 8 sets) +/- SE. Germination was defined as cotyledon emergence from the seed coat. In panel A, closed circles represent individual data points.

**Table S1.** Primers used in study

| **Gene** | **Primer orientation / name** | **Sequence** |
|---|---|---|
| **primer pair for testing CTPS splicing** | | |
| *CTPS2* (*AT3G12670*) cDNA | For. / JP1005 | AGCATGTGGTCTTCGTGTCA |
| | Rev. / WLO1400 | TAGTCCCGCCTAATTCGATG |
| *ACTIN2* (*AT3g18780*) cDNA | For. / WLO1401 | GGCTGAGGCTGATGATATTC |
| | Rev. / WLO1402 | TCTGTGAACGATTCCTGGAC |
| **qPCR primer pairs** | | |
| *ACTIN2 At3g18780* | For. / JP199 | GCACTTGCACCAAGCAGCAT |
| | Rev. / JP200 | CCTTTCAGGTGGTGCAACGAC |
| *LHCB 1.2, CAB3 At1g29910* | For. / JP197 | GGACTTGCTTTACCCCGGTG |
| | Rev. / JP198 | TCGGTAGCAAGACCCAATGG |
| *LHCB 2.2, At2g05070* | Rev. / WLO1529 | GCTTTGTAAACTCGTGATTGTG |
| | For. / WLO1530 | TGCCAAATTCACATCAAACG |
| *RBCS 2B, At5G38420* | Rev. / WLO1448 | GCTTCACCGAAGCTTAATCC |
| | For. / WLO1449 | CCACATAGAAATGGGTTCCAG |
| *CA1 AT3g01500* | For. / JP209 | TGTGTCCATCACACGTTCTGG |
| | Rev. / JP210 | GGACCACGAAGGCATCTCCT |
| *psaJ AtCG00630* | Rev. / WLO1539 | GTACTCTATGGTTCGGTTCGT |
| | For. / WLO1540 | AGGGAAATGTTAATGCATCTGG |
| *psbA AtCG00020* | For. / WLO1535 | CGTCTTTACATTGGATGGTTTGG |
| | Rev. / WLO1536 | CAGAAGTTGCGGTCAATAAGG |
| *psbB AtCG00680* | For. / WLO1537 | TCGTGCGACTTTGAAATCTGA |
| | Rev. / WLO1538 | CAACCTCTTGGGCTGCTACG |
| *rbcL AtCG00490* | For. / WLO1541 | TTACAAAGGACGATGCTACCACAT |
| | Rev. / WLO1542 | TGAGTTTCTTCTCCTGGAACGG |
| *clpP AtCG00670* | For. / WLO1545 | TGGGTTGACATATACAACCGACTTT |
| | Rev. / WLO1546 | GCCTAAAAAAAATAATCTTTCTCGATAAA |
| *rpoB AtCG00190* | For. / WLO1547 | ATACGAGATATCCATCCTAGTCAC |
| | Rev. / WLO1548 | GTCCAACATTGATTCCTTCAGAC |
| *trnEYD ATCG00250-230* | For. / WLO1559 | TCTAGTGGTTCAGGACATCTC |
| | Rev. / WLO1560 | TTGCCAACGAATTTACAGTC |
| *accD ATCG00500* | For. / WLO1563 | TTATTGCCGAACCCTATGCC |
| | Rev. / WLO1564 | AGATTCAGCCGCTTGTGAAC |
| *SIB1 At3g56710* | For. / JP589 | CAACCGGAGCCCATCTATT |
| | Rev. / JP590 | GGAGAAAGGTTGTGGTCGTC |
| *HSP26.5 At1g52560* | For. / JP585 | CGAGCTTATCGTTGCCTGAT |
| | Rev. / JP586 | CTCCGCCTTAATGTCCTCAA |
| *BAP1 At3g61190* | For. / JP338 | ATTGATGGATACGGTGGCCG |
| | Rev. / JP339 | CAGACCCCAAACCGGAACTC |
| *Atpase At3g28580* | For. / JP336 | GAAGATCGGAAAAGCGTGGAA |
| | Rev. / JP337 | CCGGGTGGTCCAAACAAAAG |

| | | |
|---|---|---|
| *ZAT12 At5g59820* | For. / JP344 | GCGTTGGTTACACGCGCTT |
| | Rev. / JP345 | CTTCAACGTAGTCACCGTGGG |
| *GST At1g17170* | For. / JP1126 | GAAGCAGCCAAGGAGTTAATCG |
| | Rev. / JP1127 | AAGCTCAGACTCTAGCGTCTTG |
| *atp1 ATMG00180* | For. / WLO1565 | GCTCCTCTGCAATTTTTGGC |
| | Rev. / WLO1566 | TGCGTGCATTCCATTATCGC |
| *ccm ATMG00180* | For. / WLO1567 | ACGTTGTTGTTTGCGAGGTC |
| | Rev. / WLO1568 | AAACATGAGCCGATCGTTCG |
| *cob1 ATMG00220* | For. / WLO1569 | TGGTTGCTTTTGGCGGATTG |
| | Rev. / WLO1570 | TCTTCCAACTCGTCCCAGAATG |
| *cox1 ATMG01360* | For. / WLO1571 | ATGCCCTTTCCAGTTTTGGC |
| | Rev. / WLO1572 | TCAGTTCAAGAGCCCAAGGAC |
| *nad1a ATMG01275* | For. / WLO1573 | ACAACCTCTAGCAGATGGTTCG |
| | Rev. / WLO1574 | ATGTAGCCACTGGAGCCATTC |
| *rpl12 ATMG00560* | For. / WLO1575 | AAGCACGTTCTCTCTTTGCG |
| | Rev. / WLO1576 | TCTTGCCGCCTAAAACTCTG |
| *CTPS1 AT1G30820* | For. / WLO1670 | ACATGGTTGAACGCCTTGAG |
| | Rev. / WLO1671 | TCCATGCGTTTGCCAGTTTC |
| *CTPS2 AT3G12670* | For. / JP873 | CTGCATTCCCAGGCATATGC |
| | Rev. / JP874 | ACGCCCGTGAAAATCAAGTC |
| *CTPS3 AT4G02120* | For. / WLO1622 | TTTTTGGCATGTCGCTCTGC |
| | Rev. / WLO1623 | ATATTAGCTGCCGCCACATG |
| *CTPS4 AT4G20320* | For. / WLO1672 | AGTTACTCAAGGGTGCTGATGG |
| | Rev. / WLO1673 | TTTCGCTGCCAGCATCTTTC |
| **Genotyping primers** | | |
| GABI-KAT Right border | Gabi-KAT 03144 | GTGGATTGATGTGATATCTCC |
| *fc2-1* (GabiKat_766H08) | LP / JP283 | GAGCAACGCCAAACATAGAAG |
| | RP / JP284 | TCAAAGGCAATGAATGTTTCC |
| *fts39/ctps2* | LP / JP770 | dCAPS markers, 146 bp PCR product, *Hpa*II restriction enzyme cuts wt sequence (121 bp) CACTCTTGGATTGCGTCAGT |
| | RP / JP771 | GACTACCTCGGAAGAACTGTACC |
| *AT3G11990* mutation in *fts39* | LP / JP768 | dCAPS markers, 126 bp PCR product, *Xba*I restriction enzyme cuts wt sequence (95 bp) GACATGGAGCTTGCTCACCA |
| | RP / JP769 | ATTAGCATCGTCTCATTTAACATGTTCTA |
| *AT3G12380* mutation in *fts39* | LP / JP836 | dCAPS markers, 121 bp PCR product, *Hpy*I88*I* restriction enzyme cuts wt sequence (99 bp) CTCTCTCCAACACTTCCTTGTT |
| | RP / JP837 | AGAAAGGGAAAAAGGGAGCAA |
| *AT3G12710* mutation in *fts39* | LP / JP859 | dCAPS markers, 132 bp PCR product, *Nco*I restriction enzyme cuts wt sequence (103 bp) AGAATCGTTGGCCACTCTCG |
| | RP / JP860 | TGAGGTGTTTAAAATCACATGTACCATG |
| *AT3G12915* mutation in *fts39* | LP / JP861 | dCAPS markers, 138 bp PCR product, *Xmn*I |

| | | |
|---|---|---|
| | | restriction enzyme cuts mutant sequence (106 bp) TGTTCACTCTTTTTACTTCATAGACCT |
| | RP / JP862 | TGAGAGGTTTATTATCGACGCTGA |
| *AT3G13175* mutation in *fts39* | LP / JP760 | dCAPS markers, 153 bp PCR product, *Eco*RI restriction enzyme cuts wt sequence (132 bp) CTTCAAAAGCTTCTCCCGGAATT |
| | RP / JP761 | agacagagagagagttagcattgt |
| **Cloning primers** | | |
| *CTPS1* (*AT1G30820*) coding region for Gateway - stop codon (3583 bp fragment of *CTPS1*) | For / WLO1392 | ggggacaagtttgtacaaaaaagcaggcttgGTAGGAAGA AGATGAAGTACGTGC |
| | Rev / WLO1393 | ggggaccactttgtacaagaaagctgggtcTCTAGTGTAG AGGCCATTGCAGTAG |
| *CTPS2* (*AT3G12670*) coding region for Gateway - stop codon (3803 bp fragment of *CTPS2*) | For / JP1192 | ggggacaagtttgtacaaaaaagcaggcttgTCTTTCCTTA CCGTGTTCTGTGT |
| | Rev / JP1193 | ggggaccactttgtacaagaaagctgggtcGTGGTGAAGC CCGTTTCCAT |
| *CTPS2* (*AT3G12670*) promoter and coding region for Gateway (7075 bp fragment including 2262 bp 5' / 1073 bp 3' of coding region) | For / JP928 | caccTGCTGACCTGTTCTGACGTC |
| | Rev / JP929 | CCAACCAACCTCCGTCATCA |
| *CTPS3* (*AT4G02120*) coding region for Gateway - stop codon (3229 bp fragment of *CTPS3*) | For / WLO1396 | ggggacaagtttgtacaaaaaagcaggcttgGGAAGAGAT GAAGTACGTATTGGTG |
| | Rev / WLO1397 | ggggaccactttgtacaagaaagctgggtcATTGCTTAAA TGGGCTTGAAGAAGC |
| *CTPS3* (*AT4G02120*) coding region for Gateway plus + stop codon (3232 bp fragment of *CTPS3* with WLO1396) | Rev / WLO 1456 | ggggaccactttgtacaagaaagctgggtcTCAATTGCTT AAATGGGCTTGAAGA |
| *CTPS4* (*AT4G20320*) coding region for Gateway - stop codon (4060 bp fragment of *CTPS4*) | For / WLO1403 | ggggacaagtttgtacaaaaaagcaggcttgTTTGCAGGA AACTTGTTAGTGTTTAAG |
| | Rev / WLO1395 | ggggaccactttgtacaagaaagctgggtcCGAGTAGACA CGATCACATAAACTG |
| *At3g11990* promoter and coding region for Gateway (3663 bp fragment including 1713 bp 5' / 1056 bp 3' of | For / JP926 | caccATCCCTGTGACCGGAAACA |

| | | |
|---|---|---|
| coding region) | | |
| | Rev / JP927 | TAGTGGCATACCGACGATGC |
| *AT3G12380* promoter and coding region for Gateway (7064 bp fragment including 2339 bp 5' / 1248 bp 3' of coding region) | For / JP922 | caccACTGCAACATCCTTACGCCT |
| | Rev / JP923 | ggtagccaactgtttcaggc |
| *AT3G12710* promoter and coding region for Gateway (4782 bp fragment including 2340 bp 5' / 1185 bp 3' of coding region) | For / JP924 | caccGGGTTGCAGAGATGTGGGAA |
| | Rev / JP925 | TGCGCTCTCTTCTCAAAGCA |
| *At3g12915* promoter and coding region for Gateway (6140 bp fragment including 2396 bp 5' / 1034 bp 3' of coding region) | For / JP930 | caccTCGATCGGTTGTGGAATTTGC |
| | Rev / JP931 | CAGCAGGCCAAGACAACAAC |
| *AT3G13175* promoter and coding region for Gateway (3609 bp fragment including 2071 bp 5' / 1217 bp 3' of coding region) | For / JP932 | caccCCGTTGTGCAAGCTGGTTTA |
| | Rev / JP933 | GGAAGAGCCAGCGAAGTCAT |

**Table S2.** Vectors used in study

| **Plasmid** | **Insert** | **notes** | **ref** |
|---|---|---|---|
| pENTR-D/TOPO | | Gateway cloning vector | Invitrogen |
| pDONR221 | | Gateway cloning vector | Invitrogen |
| PEARLEYGATE-101 | | 35S promoter, C-term YFP-HA fusion | (Earley *et al.*, 2006) |
| PEARLEYGATE-102 | | 35S promoter, C-term CFP-HA fusion | (Earley *et al.*, 2006) |
| pGBGWY | | No promoter, C-term YFP | (Zhong *et al.*, 2008) |
| pWLP314 | *CTPS1* coding region (gDNA) – stop codon | pDONR221 | This study |
| pJDW224 | *CTPS2* coding region (gDNA) – stop codon | pDONR221 | This study |
| pWLP334 | Mut *CTPS2* coding region (gDNA) – stop codon | pDONR221 | This study |

| | | | |
|---|---|---|---|
| pJDW174 | Genomic fragment of *CTPS2* coding region plus 2262 bp 5' / 1073 bp 3' | pENTR-D/TOPO | This study |
| pWLP316 | *CTPS3* coding region (gDNA) – stop codon | pDONR221 | This study |
| pWLP348 | *CTPS3* coding region (gDNA) + stop codon | pDONR221 | This study |
| pWLP315 | *CTPS4* coding region (gDNA) – stop codon | pDONR221 | This study |
| pWLP322 | *35S::CTPS1-YFP-HA* | In PEARLEYGATE-101 | This study |
| pCTPsyn1 | *UBQ10::CTPS1*(cDNA)-*YFP* | pHygII_UT_mVenusC (Kunz *et al.*, 2014) | (Daumann *et al.*, 2018) |
| pJDW229 | *35S::CTPS2-YFP-HA* | In PEARLEYGATE-101 | This study |
| pWLP345 | *35S::mutCTPS2-YFP-HA* | In PEARLEYGATE-101 | This study |
| pJDW181 | *CTPS2::CTPS2* | pGBGWY | This study |
| pCTPsyn2 | *UBQ10::CTPS2*(cDNA)-*YFP* | pHygII_UT_mVenusC (Kunz *et al.*, 2014) | (Daumann *et al.*, 2018) |
| pWLP324 | *35S::CTPS3-YFP-HA* | In PEARLEYGATE-101 | This study |
| pWLP351 | *35S::CTPS3-CFP-HA* | In PEARLEYGATE-102 | This study |
| pWLP349 | *35S::CTPS3* (native) | In PEARLEYGATE-101 | This study |
| pCTPsyn3 | *UBQ10::CTPS3*(cDNA)-*YFP* | pHygII_UT_mVenusC (Kunz *et al.*, 2014) | (Daumann *et al.*, 2018) |
| pWLP323 | *35S::CTPS4-YFP-HA* | In PEARLEYGATE-101 | This study |
| pWLP350 | *35S::CTPS4-CFP-HA* | In PEARLEYGATE-102 | This study |
| pCTPsyn4 | *UBQ10::CTPS4*(cDNA)-*YFP* | pHygII_UT_mVenusC (Kunz *et al.*, 2014) | (Daumann *et al.*, 2018) |
| pCTPsyn5 | *UBQ10::CTPS5-YFP* | pHygII_UT_mVenusC (Kunz *et al.*, 2014) | (Daumann *et al.*, 2018) |
| pJDW177 | *AT3G12380*:: | pGBGWY | This study |

| | | | |
|---|---|---|---|
| | *AT3G12380* | | |
| pJDW178 | *AT3G12710*:: *AT3G12710* | pGBGWY | This study |
| pJDW179 | *AT3G11990*:: *AT3G11990* | pGBGWY | This study |
| pJDW180 | *AT3G13175*:: *AT3G13175* | pGBGWY | This study |
| pJDW182 | *AT3G12915*:: *AT3G12915* | pGBGWY | This study |

**Table S4.** Complementation and phenocopy analyses of overexpression vectors

| **Overexpressed gene** | **Construct** | **Genetic background** | **Complement *ctps2*? (# T1's / total T1's)** |
|---|---|---|---|
| *none* | Empty *pEARLEY101* | *fc2-1 ctps2* | 0/19 |
| *CTPS1 At1g30820* | *35S::CTPS1-YFP-HA* | *fc2-1 ctps2* | 11/14 |
| *CTPS1 At1g30820* | *UBQ10::CTPS1-YFP* | *fc2-1 ctps2* | 4/6 |
| *CTPS2 At3g12670* | *35S::CTPS2-YFP-HA* | *fc2-1 ctps2* | 10/10 |
| *CTPS2 At3g12670* | *UBQ10::CTPS2-YFP* | *fc2-1 ctps2* | 18/21(Zhong *et al.*, 2008)(Zhong *et al.*, 2008) |
| *CTPS3 At4g02120* | *35S::CTPS3-YFP-HA* | *fc2-1 ctps2* | 0/17* |
| *CTPS3 At4g02120* | *UBQ10::CTPS3-YFP* | *fc2-1 ctps2* | 0/4* |
| *CTPS3 At4g02120* | *35S::CTPS3* | *fc2-1 ctps2* | 0/25 |
| *CTPS4 At4g20320* | *35S::CTPS4-YFP-HA* | *fc2-1 ctps2* | 17/20 |
| *CTPS4 At4g20320* | *UBQ10::CTPS4-YFP* | *fc2-1 ctps2* | 7/8 |
| *CTPS5 At2g34890* | *UBQ10::CTPS5-YFP* | *fc2-1 ctps2* | 5/7 |
| **Overexpressed gene** | **Construct** | **Genetic background** | **phenocopy *ctps2*? (# T1's / total T1's)** |
| *mutCTPS2 At3g12670* | *35S::mutCTPS2-YFP-HA* | *fc2-1* | 0/9* |

* only include lines with observable YFP levels

**Table S5.** *ctps* mutant alleles

| ***ctps* allele** | **Original name** | **Type of mutation** | **ref** |
|---|---|---|---|
| ctps2-1 | *Gabi-Kat_032C02* | intron 14 | (Daumann *et al.*, 2018) |
| ctps2-2 | *Gabi-Kat_156G07* | intron 6 | (Daumann *et al.*, 2018) |
| ctps2-3 | *emb2742-1* | intron 5 | (Meinke, 2020) |
| ctps2-4 | *emb2742-2* | promoter region | (Meinke, 2020) |

| | | | |
|---|---|---|---|
| ctps2-5 | *fts39* | G840A, splicing mutant | This study |
